# A human lysosomal storage disorder toolkit for decoding proteome landscapes in cortical and dopaminergic-like induced neurons

**DOI:** 10.1101/2025.10.08.681047

**Authors:** Felix Kraus, Yuchen He, Yizhi Jiang, Delong Li, Yohannes A. Ambaw, Federico M. Gasparoli, Joao A. Paulo, Tobias C. Walther, Robert V. Farese, Steven P. Gygi, Florian Wilfling, J. Wade Harper

**Affiliations:** Department of Cell Biology, Harvard Medical School, Boston MA, USA; Aligning Science Across Parkinson’s (ASAP) Collaborative Research Network, Chevy Chase, MD 20815, USA; Mechanisms of Cellular Quality Control, Max Planck Institute of Biophysics, Frankfurt, Germany; Cell Biology Program, Sloan Kettering Institute, New York, NY, USA; Howard Hughes Medical Institute, New York, NY, USA

**Author notes:** Corresponding Author: J. Wade Harper, **Email:**. Department of Pathology, University of British Columbia, Vancouver, BC, Canada. **Author Contributions:** Conceptualization: F.K., J.W.H.; Investigation: F.K., Y.H., D.L., J.A.P., Y.J., Y.A.A; Analysis: F.K., D.L., F.M.G., J.W.H.; Visualization: F.K. Supervision: R.V.F., T.C.W., S.P.G., F.W., J.W.H.; Writing (original draft): F.K., J.W.H.; Writing (reviewing and editing): F.K., J.W.H., with input from all authors. **Competing Interest Statement:** J.W.H. is a co-founder of Caraway Therapeutics, a subsidiary of Merck & Co., Inc., Rahway, NJ, USA (Caraway) and is a scientific advisory board member for Lyterian Therapeutics. This publication is unrelated to these competing interests. All other authors have no competing interests to declare.

**Keywords:** Proteomics, Organelle quality control, iNeuron, iDA, Lysosomal Storage Disorder

## Abstract

Lysosomes maintain cellular homeostasis by degrading proteins delivered via endocytosis and autophagy, and by recycling building blocks for organelle biogenesis. Lysosomal Storage Disorders (LSDs) comprise a group of diseases affecting diverse lysosomal functions. To facilitate molecular phenotyping across diverse LSD gene classes, we are developing a library of human embryonic stem cells engineered to lack individual LSD genes as a resource for the field. Here, we report our initial stem cell toolkit lacking one of 23 LSD genes, including the majority of genes associated with sphingolipidoses and neuronal ceroid lipofuscinoses, and its use in the generation of a proteomic resource for induced cortical-like and midbrain dopaminergic-like neurons In-depth abundance and correlation profiling across organelles and sub-organelle components revealed potential vulnerabilities that reflect distinct patterns of proteome alterations across both genotypes and neuronal cell types. We characterize alterations in the mitochondrial proteome associated with GBA1 and ASAH1 deficiency and identify synaptic and mitochondrial defects in *ASAH1*^-/-^ induced neurons that correlate with defects in neuronal firing rates. Moreover, we developed an informatic pipeline for proteome-wide identification of individual protein interactions and protein complexes that may be disrupted as a result of LSD gene deficiency. Finally, we visualized structural alterations of *ASAH1*-deficient endolysosomes *in situ* using cryo-electron tomography, revealing swollen organelles that were largely devoid of dense internal membranes characteristic of wild-type cells, but containing numerous intralumenal vesicle compartments. This toolkit and associated proteomic landscapes provide a resource for defining molecular signatures associated with LSD gene dysfunction and organelle vulnerability.

## Introduction

Lysosomes are a primary degradative membrane-bound compartment within cells, and contain hydrolases that digest glycoproteins and lipids delivered via endocytosis, as well as proteins and organelles delivered via autophagy, thereby facilitating the recycling of molecular building blocks(1, 2). Additionally, lysosomes contain transporters responsible for trafficking of recycled building blocks to other cellular compartments. As such, lysosomes function as important signaling hubs for nutrient availability via the MTOR pathway (1). Genetic and clinical studies have identified ∼70 distinct types of lysosomal storage disorders (LSDs) that map to at least 54 genes directly or indirectly linked with lumenal catabolic functions. LSDs are primarily classified as sphingolipidoses, mucopolysaccharidoses, glycoproteinoses, and neuronal ceroid lipofuscinoses (NCL), based on the type of storage material that accumulates within lysosomes (e.g. glycosphingolipids, or cholesterol) (3). Additional storage material can also accumulate as a secondary response. However, how diverse primary and secondary storage material accumulation interfaces with distinct multisystem disease phenotypes is poorly understood. One possibility is that various LSD risk alleles create molecular vulnerabilities that can be mapped onto organelles and pathways in a cell-type dependent manner, potentially providing a deeper understanding of LSD pathogenesis mechanisms. In this context, “vulnerabilities” refer to alterations in protein abundance that can increase or decrease that may be inferred to affect a protein’s overall cellular activity or that of its associated protein complex. Such alterations in proteins that are central to organelle function or associated quality control pathways could adversely affect organelle function. As such, landscapes of protein vulnerabilities, integrated across organelles and key protein complexes, provide starting points toward a systematic understanding of how disease risk alleles impact organelle and cell systems.

As an initial approach to addressing this knowledge gap, we previously reported a library of more than 2 dozen HeLa cell lines lacking genes linked with LSDs, facilitating global lipidomic and proteomic profiling and revealing patterns of alterations in autophagy and cholesterol trafficking systems with specific LSD mutations (4). Here, we extend this approach through the development of a versatile toolkit of human embryonic stem (ES) cell lines lacking one of 23 LSD genes, facilitating the generation of a resource for proteomic profiling of cortical-like and midbrain dopaminergic-like induced neuron (iN and iDA) models across LSD alleles. The current toolkit is focused on genes linked with sphingolipidoses, NCL, and integral membrane protein disorder classes. The focus on sphingolipidoses and NCL genes reflects: 1) an emerging framework for understanding the mechanisms and phenotypes associated with defects in sialic-acid-containing glycosphingolipid breakdown on intraluminal vesicles (ILVs) formed within endolysosomes, including the role of bis(monoacylglycerol)phosphate (BMP) lipids in this process (5-8), 2) the proposed genetic linkage between glycosphingolipid breakdown enzymes GBA1, SMPD1, ASAH1 and Parkinson’s disease (PD) (9-18), and 3) the finding that specific NCL genes, including CLN11 (also called GRN), may regulate the abundance of BMP within lysosomes, with loss of BMP resulting in the accumulation of GM2 and GM3 gangliosides (7). Using deep global proteome analysis of day 50 iN and iDA cells coupled with informatic pipelines for decoding organelle protein abundance landscapes and underlying protein-protein interaction (PPI) networks, we identify patterns of proteome alterations that may define functional vulnerabilities associated with specific LSD genes. Candidate vulnerabilities include alterations in mitochondrial proteomes particularly in *GBA1*^-/-^ iDA cells, as previously observed (19, 20), and synaptic proteomes in *ASAH1*^-/-^ cells, with *ASAH1*^-/-^ iDA cells displaying reduced neuronal firing. Finally, to understand the impact of ASAH1 deficiency on lysosomal ultrastructure, we performed cryo-electron tomography (cryo-ET) on lysosomal structures in the HeLa cell system, revealing swollen structures largely devoid of lamellar membrane material typically found in lysosomal structures in wild-type HeLa cells. This toolkit and proteomic data made available through an online viewer provides a resource for the field.

## Results

### A Stem Cell Toolkit for Systematic Analysis of LSD Genes

To systematically examine potential alterations in the properties of organellar systems in the context of LSDs, we developed a versatile stem cell loss of function toolkit. We employed an H9 human embryonic stem cell (hESC)-based system (4, 21) containing an inducible AAVS1-NGN2 cassette for efficient neuronal differentiation and endogenously tagged TMEM192-3xHA (heterozygous) for LysoIP (H9^NGN2;TMEM192-HA^). These cells, referred to hereafter as “Control” cells, can be efficiently convert to cortical-like iNeurons (iN) or midbrain-like dopaminergic neurons (iDA) using established methods (22, 23), as demonstrated here by immunofluorescence with α-MAP2 and α-TH (Tyrosine Hydroxylase) (***SI Appendix*, Fig. S1*A***). The initial version of this toolkit includes inactivating indels in 23 LSD genes, including genes implicated in sphingolipidoses (*GBA1, ASAH1, HEXA, HEXB, PSAP, SMPD1*), NCL (*CLN1, CLN2, CLN3, DNAJC5, CLN5, CLN6, CLN7, CLN8, ATP13A2, GRN, CTSD, CTSF*), Mucolipidosis type IV (*MCOLN1* (encoding TRPML1), as well as Niemann-Pick Type-C (*NPC1, NPC2*), *LIPA* (Wolman’s disease) and *GAA* (Pompe’s disease). CRISPR RNPs were transfected into Control cells and homozygous clones with frameshift indels were identified by locus sequencing (see **Dataset S1** for a detailed sequence analysis of indels and clone identification). Proteomic data described in the following section confirms loss of function mutations, and points to the utility of this system for our ongoing effort to create a complete LSD toolkit.

### Proteomic Fingerprint Resource for LSD Mutant iN and iDA Neurons

Our previous work has indicated the utility of total proteome analysis to reveal alterations in organelle and pathway abundance in response to loss of specific pathways (e.g. autophagy) or during changes in cell state (24-26). To profile alterations in the proteome of our LSD toolkit, we prioritized generation of day 50 iN and iDA cells in biological triplicate for 23 individual mutant lines along with Control cells (**Fig. 1*A* and Dataset S2**), which is a sufficient period for creation of functional synapses (22, 23). In parallel, a focused set of cell lines lacking genes directly (*ASAH1, GBA1, SMPD1*) or indirectly (*GRN*) (7) linked with sphingolipidoses were differentiated for 30 days, in biological triplicates (**Fig. 1*A* and Dataset S3**). Total cell proteins and interspersed quality control samples were subjected to analysis using an Astral mass spectrometer with nDIA-hrMS2 (narrow-window data-independent acquisition-high resolution MS^2^) acquisition (**Fig. 1*A***), allowing detection of ∼10,000 proteins across cells of each genotype (***SI Appendix*, Fig. S1*C***,***D***). This 30 day experiment allowed for an interim quality control and proteomic analysis at an intermediate stage of differentiation while longer term cultures continued differentiation, while also facilitating an analysis of continued differentiation through neuronal marker analysis. Importantly, we observed consistent instrument performance over extended data collection periods (standard deviation <10%) for both the 30 day and 50 day differentiation periods (***SI Appendix*, Fig. S1*E*** and **Dataset S2**,**3**), pointing to the consistency and robustness of the dataset.

**Fig. 1.**
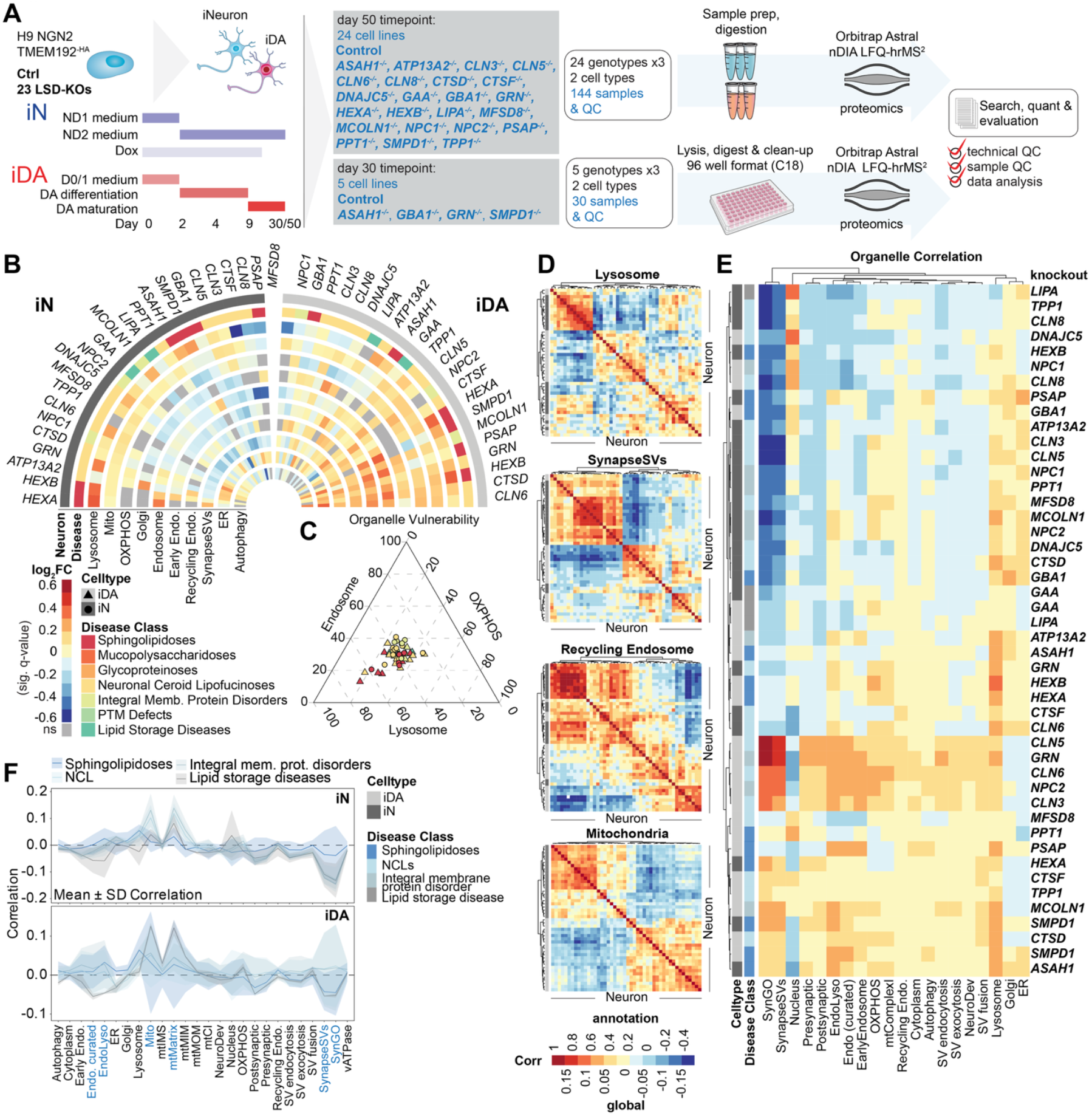
Proteome atlas of lysosomal Storage Disorder neuronal cell models. **(*A*)** Schematic of experimental workflow for proteomic resource generation. Control and 23 LSD knockout human ES cell lines were differentiated to iN and iDA cells for 50 days in biological triplicate and analyzed using nDIA LFQ. Independently, four LSD mutant cell lines and Control cells were differentiated for 30 days followed by proteomic analysis. **(*B*)** Half-circos plot of log_2_FC mean protein abundance per annotation (q < 0.05, FDR-corrected) of each LSD knockout/Control for iN (left) and iDA (right) neurons for the indicated annotations or organelles. **(*C*)** Ternary plot of log_2_FC [KO/Control] for endosome, lysosome and OXPHOS annotations across iN and iDA neurons reveals clusters of genotypes with altered endolysosomal proteomes. Triangles, iDA cells; circles, iN cells. **(*D*)** Example of pairwise, proteome-wide cross-correlation of lysosome, SynapseSV (middle), recycling endosome and mitochondria annotated proteins (from top to bottom). **(*E*)** Composite correlation across 20 organelle annotations. **(*F*)** Mean organelle correlation for the four major disease classes (Sphingolipidoses, NCLs, Integral membrane protein disorders, and Lipid storage disorders). ± SD for iN (top) and iDA (bottom). Disease class is indicated by shading.

We validated the robustness of the dataset as a resource by performing multiple types of quality control analyses. First, we compared the proteomes of Control iN and iDA cells. We detected 8,920 proteins with comparative relative abundance in both cell types with ∼500 proteins selectively enriched (log_2_FC>1.0, p-value<0.05) in either iN or iDA cells (***SI Appendix*, Fig. S1*F* and Dataset S2**,**3**). These differentially altered proteins were distributed across multiple subcellular compartments with the largest number of differentially enriched proteins present in the nucleus or synaptic compartments (***SI Appendix*, Fig. S1*G***). The 104 selectively enriched proteins annotated within the SynapseSV group (see **Dataset S1**) were distributed in iDA and iN cells (∼41% and ∼61%, respectively, with the highest Gene Ontology (GO) categories related to synaptic vesicle transport (***SI Appendix*, Fig. S1*H***). In contrast, the mitochondrial compartment was skewed towards enrichment in iDA cells (∼86% versus ∼14% in iN cells), with oxidative phosphorylation represented prominently in the GO analysis (***SI Appendix*, Fig. S1*H***). Consistent with the differentiation of iDA cells, TH was found to be enriched in this population in our unbiased analysis (**Dataset S2**), with MS1 intensity being ∼6-fold higher in iDA than iN cells (***SI Appendix*, Fig. S1*I***). Likewise, SYN1 and NEFH were also identified as iDA-enriched proteins, and differential expression was validated using immunofluorescence of day 38 iDA or iN cells (***SI Appendix*, Fig. S1*J***). In contrast with TH, the potassium channel protein KCNJ6 did not reach the 2-fold threshold for enrichment in iDA cells (***SI Appendix*, Fig. S1*I***). As expected, several neuronal differentiation markers detected in 30 day Control iN or iDA cells displayed further increase in abundance at 50 days of differentiation (***SI Appendix*, Fig. S1*K***), with a similar trend seen with groups of presynaptic, postsynaptic, and “neurondriver” proteins (***SI Appendix*, Fig. S1*L***, see **METHODS** for list of neuronal and synaptic marker proteins used in this analysis).

We next performed a series of quality control experiments across the LSD mutant cell lines. First, we found that Pearson coefficients of pairwise replicates were typically >0.98 (***SI Appendix*, Fig. S2*A***). Second, principal component (PC) analysis revealed separation of neuronal subtype and length of differentiation (***SI Appendix*, Fig. S2*B***). Third, we observed the expected loss of specific LSD proteins based on proteomics, although we note that CLN3, CLN8 and MSFD8 proteins were not detected at the whole cell level and likely require enrichment for detection (***SI Appendix*, Fig. S2*C; Dataset S1***). Fourth, the various LSD mutant cells demonstrated the expected induction of several neuronal markers by proteomics (***SI Appendix*, Fig. S2*D***, see METHODS for list of neuronal marker proteins used in this analysis), and TH was consistently expressed across all iDA cells (***SI Appendix*, Fig. S2*E***).Taken together, our data supports the utility of this cell line and proteomic resource for further exploration both of iN and iDA lineages and the impact of LSD protein deletion.

### LSD Mutant iNeuron Organelle Landscapes

We initially examined alterations in protein abundance, finding that the number of proteins significantly changed compared to Control (q-value < 0.05, FDR-corrected) varied with genotype and cell type (***SI Appendix*, Fig. S3*A***). This reflected distinct patterns of alterations in the abundance of specific organelles or protein complexes/pathways (e.g. autophagy) (**Fig. 1*B***). Alterations in the abundance of proteins assigned to individual organelles provides a means by which to explore possible organelle vulnerability across genotype and disease class, with, for example, a subset of iDA and iN cells of sphingolipidoses and NCL cells displaying the largest alterations of changes in protein abundance for lysosome and endosome compartments (**Fig. 1*C***). The ability to identify organelle vulnerability signatures within the proteomic data depends upon the quality of the underlying annotations. For example, although various synaptic protein categories in the GO database are contained within SYNGO, there are varying degrees of overlap within the various GO groups (***SI Appendix*, Fig. S3*B***). We therefore used these more refined compartment annotations along with an analogous treatment of the endolysosomal compartment (***SI Appendix*, Fig. S3*B* and Dataset S1**) to perform hierarchical clustering of protein log_2_FC alterations across organelle and sub-cellular annotations (***SI Appendix*, Fig. S3*C***). This analysis revealed shared patterns of increased or reduced protein abundance especially for synaptic vesicle, lysosomal, and mitochondrial annotations, with iDA and iN cell types sometimes demonstrating distinct patterns (***SI Appendix*, Fig. S3*C***). F-ratio analysis of genotype variance by annotation (between vs. within) demonstrated that the most impacted organelle annotations reflected biological signals rather than technical noise (F-ratio>1.0) (***SI Appendix*, Fig. S3*D***).

To capture the global impact of each LSD gene deletion, we generated an organelle correlation map by computing all pairwise correlations of annotation-specific mean log_2_fold-changes across genotypes and cell types (**Fig. 1*D,E***). This composite map integrates the direction and strength of these relationships (as a correlation factor) to reveal how each knockout reshapes proteome organization across subcellular compartments. For example, both *CLN5*^*-/-*^ and *GRN*^*-/-*^ iDA cells share high similarity in synaptic annotation categories (SynGO, SynapaseSVs), whereas *SMPD1*^*-/-*^ and *ASAH1*^*-/-*^ iN cells of clustered together with *SMPD1*^*-/-*^ and *CTSD*^*-/-*^ iDA cells (but interestingly not ASAH1^*-/-*^ iDA) (**Fig. 1*E***). The highest variability in correlations across disease classes was found for groups of proteins linked with mitochondria, endolysosome, and synaptic vesicles/SynGO, for both iN and iDA neurons (**Fig. 1*F***). For example, the SynapseSV category of proteins displays high variance across genotypes, particularly in iDA neurons, while in iN cells, the sphingolipidoses group of proteins displayed the most extensive variability (**Fig. 1*F***). Distinct patterns of variation between organelles for the two cell types can be visualized in correlation plots for particular LSD mutations (**Fig. 2*A***). For example, synaptic proteins are among the most variable (most negative correlations) in both iDA and iN *GBA1*^-/-^ (Sphingolipidoses) cells, while mitochondria are selectively affected (negatively correlated) in iDA cells (**Fig. 2*A***). In contrast, synaptic proteins were positively correlated in iDA *GRN*^-/-^ (NCL) or *MCOLN1*^-/-^ cells, but were negatively correlated in iN cells (**Fig. 2*A***). As an orthogonal approach to comparing the effects of LSD mutations, we implemented a ranked impact score that assesses the extent of variability in subcellular compartments in either iDA or iN cells of individual disease classes, with the lowest score reflecting the greatest impact (**Fig. 2*B***) (see ***SI Appendix*, MATERIALS and METHODS**). Taken together, this data suggests a complex interplay between genotype and cell type in defining organelle landscapes across LSDs, and led us to focus further on mitochondrial and synaptic compartments.

**Fig. 2.**
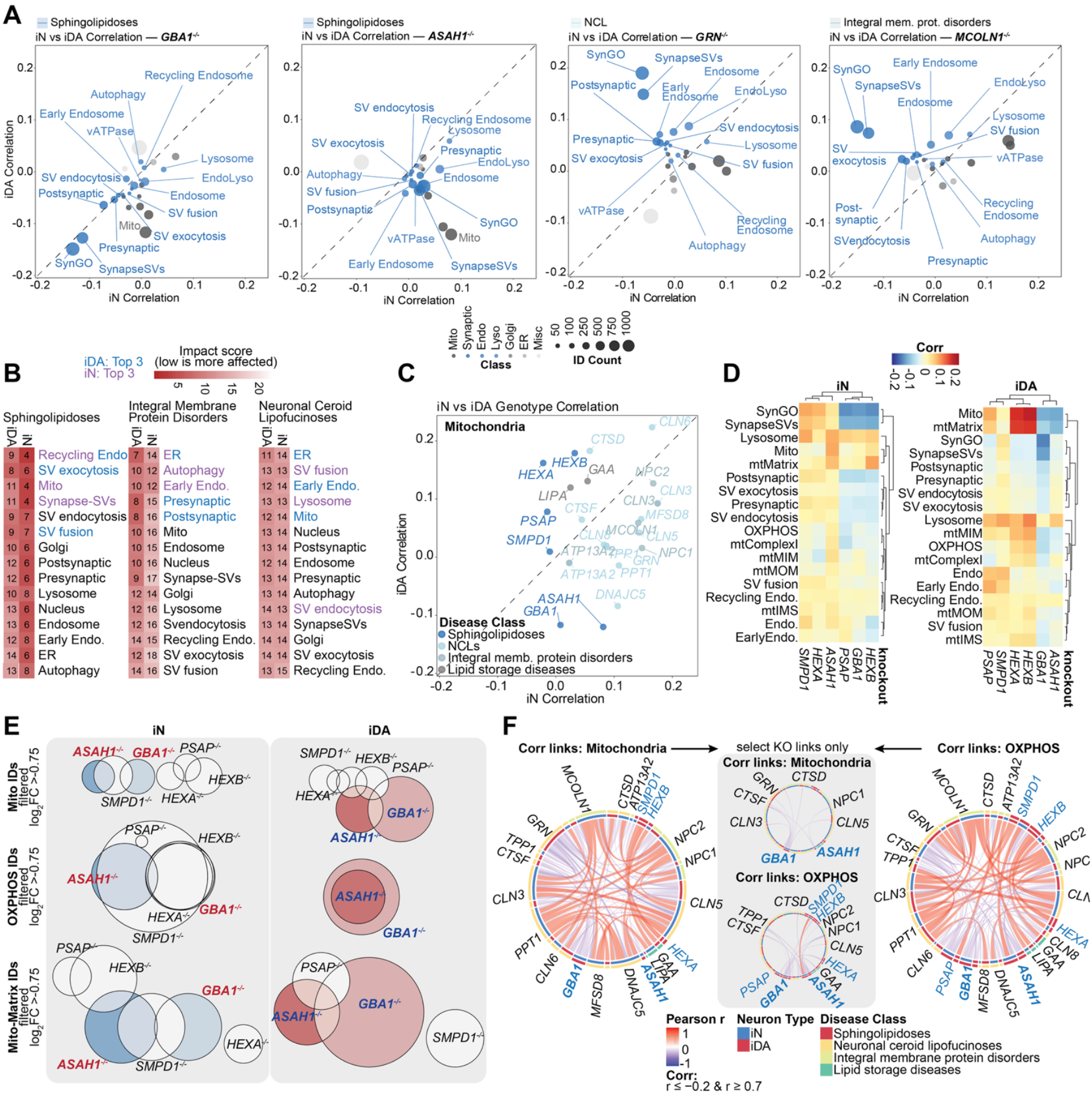
Differential mitochondrial organelle vulnerability across Sphingolipidoses mutant iDA neurons. **(A)** iN vs. iDA organelle and protein complex/compartment correlation plots for cells deficient in *GBA1, ASAH1, GRN*, or *MCOLN1* (TRPML1). **(*B*)** Impact score reflecting the extent to which genotypes within Sphingolipidoses, Integral Membrane Protein Disorders, and Neuronal Ceroid Lipofuscinoses categories display alterations across various organelles and protein complex/compartment proteomes in either iN or iDA cells. The numbers in each square represent the rank in each cell lineage. The lower the number indicates a more severe disruption across the genotypes examined. **(*C*)** iN vs. iDA correlation plot for mitochondria across 23 LSD mutants. **(*D*)** Hierarchical cluster correlation heatmap of indicated organelle annotations across Sphingolipidoses mutants for iN (left) and iDA (right) cells. **(*E*)** Euler plots depicting overlap and unions of mitochondrial (top), OXPHOS (middle) and mito-matrix (bottom) protein IDs log_2_FC > -0.75 across sphingolipidoses mutants for iN and iDA cells. **(*F*)** Circos plots showing correlations between genotypes for mitochondria (left) and OXPHOS (right). Center panel depicts genotypic links originating from only *GBA1*^*-/-*^ and *ASAH1*^*-/-*^. Only links with Pearson scores (r) smaller than -0.2 and larger than 0.7 are shown. Blue labels indicate genotypes belonging to Sphingolipidoses disease class. Bold labels are PD-risk genes.

### Sphingolipidoses-related Mitochondrial Proteome Signatures

Previous linkage of *GBA1* mutants with mitochondrial dysfunction (19, 20) together with mitochondria being among the most impacted organelles with Sphingolipidoses-related mutants (**Fig. 2*A***) led us to examine this relationship in detail. Correlation plots for iN and iDA cells for the mitochondrial proteome revealed extensive variation for Sphingolipidoses mutants in iDA cells but much less so in iN cells (**Fig. 2*C***). Interestingly, while both *GBA*1^-/-^ and *ASAH1*^-/-^ iDA cells displayed negative correlations, *HEXA*^-/-^, *HEXB*^-/-^ and *PSAP*^-/-^ positive correlations (**Fig. 2*C***), indicative of distinct behavior, which was further visualized through unsupervised cluster analysis of individual organelle annotations (**Fig. 2*D***). We separated specific mitochondrial proteins based on functional categories and abundance, including proteins involved in oxidative phosphorylation (OXPHOS) and mitochondrial matrix (mtMatrix), and examined overlap between the various Sphingolipidoses genotypes in iN and iDA cells (**Fig. 2*E***). While *GBA1*^-/-^ and *ASAH1*^-/-^ mutants displayed either complete (OXPHOS) or partial (mito- and mtMatrix) overlap in proteins with altered abundance in iDA cells, no overlap was observed in the context of iN cells (**Fig. 2*E***). Consistent with OXPHOS components underlying the dependency on neuron subtype, negative correlation connectivity (links) with other LSD gene deletions were prominent for *ASAH1*^-/-^ iDA cells in the context of OXPHOS proteins, but were much less prominent in the context of total mitochondrial proteins (**Fig. 2*F***). Interestingly, the only positive link observable in the dataset for *ASAH1*^-/-^ cells was with *SMPD1*^*-/-*^, and this was specific to iN cells (**Fig. 2*F***). Together, these data provide insight into how loss of specific Sphingolipidoses genes alters the abundance of selective classes of proteins within the mitochondrial compartment in a neuronal type-dependent manner.

### Sphingolipidoses-related Synaptic Proteome Signatures

Impact scores for synaptic and endocytic components among Sphingolipidoses mutants (**Fig. 2*B***) led us to focus on alterations in these compartments. Cluster analysis of log_2_FC data for individual organelles revealed distinct patterns across the LSD genotypes (**Fig. 3*A,B***). For example, some genotypes, including all Sphingolipidoses mutants, displayed increased levels of lysosomal proteins, which was frequently mirrored by reduced levels of synaptic proteins. In both cases, *ASAH1*^-/-^ mutants displayed a small negative correlation for SynapseSV and SV fusion annotations in iDA cells but a positive correlation in iN cells, suggesting a distinct behavior (**Fig. 3*C,D***). Most Sphingolipidoses cells displayed reductions in SV endocytosis and SV exocytosis protein abundance, but unlike other genotypes, *GBA1*^-/-^ and *ASAH1*^-/-^ cells also showed reductions in the abundance of SV fusion proteins selectively in iDA neurons (**Fig. 3*B***). Proteins within the SV fusion category displayed numerous negatively correlated links with specific LSD mutants in either iDA or iN cells, and as expected, the number of such links within the SynapseSV category was substantially reduced relative to SV fusion (**Fig. 3*E***). These data suggest potential synaptic vulnerability in *ASAH*^-/-^ iNeurons, which is address below.

**Fig. 3.**
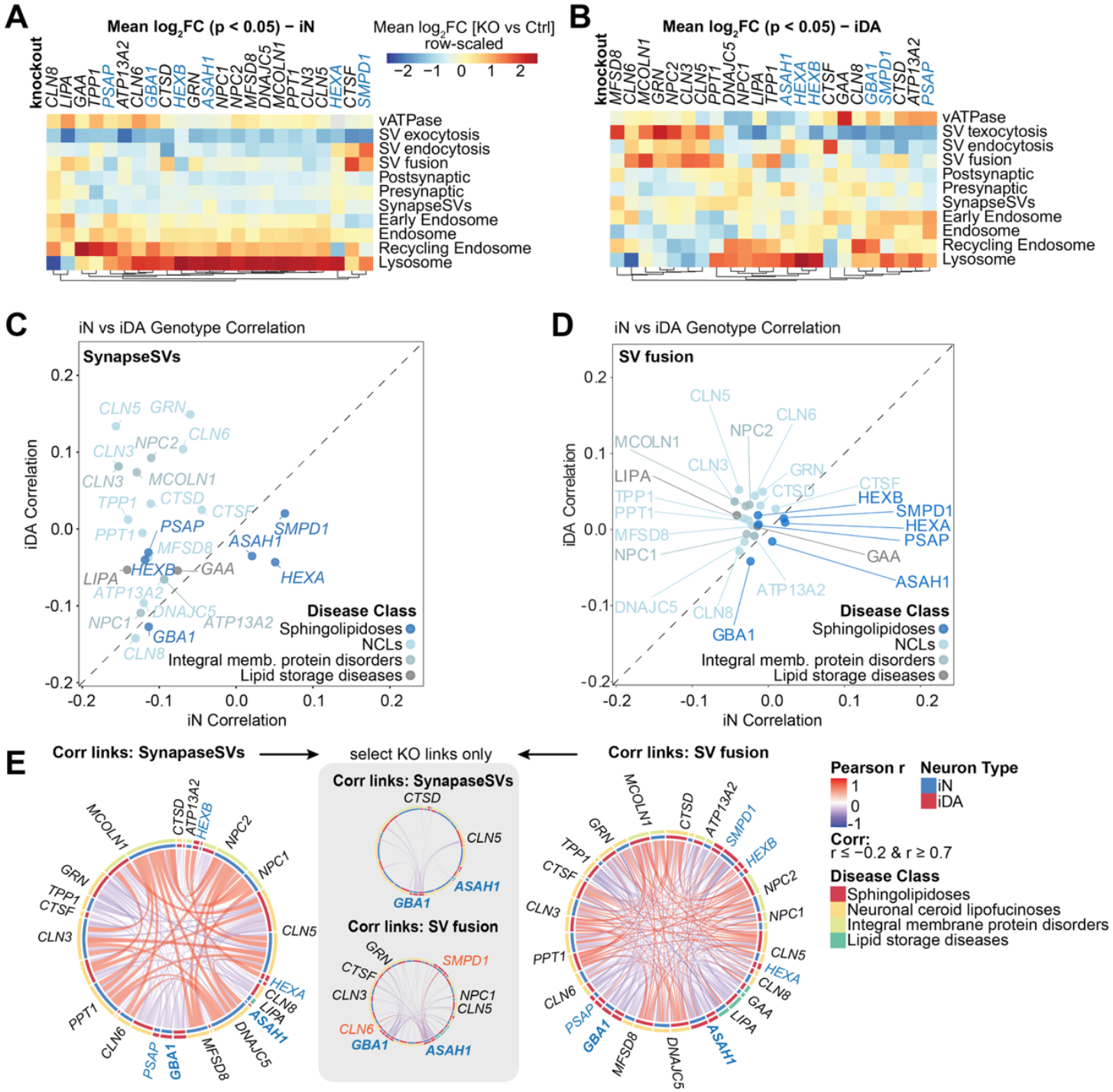
Systematic analysis of synaptic and endolysosomal compartments across LSD iN and iDA cells. **(*A***,***B*)** Log_2_FC [KO vs Control] heatmap of vesicular annotations for iN (panel ***A***) and iDA (panel ***B***) cells at day 50 of differentiation. The annotations are sorted from high to low from left to right based on iN cell data. Sphingolipidoses related genotypes are in blue and LSD-PD risk genes are labeled in bold. **(*C***,***D*)** iN vs. iDA organelle correlation of SynapseSV (panel ***C***) and SV fusion (panel ***D***) proteins. Genotypes are color-coded by disease class. **(*E*)** Circos plots showing correlations between genotypes for SynapseSV (left) and SV fusion (right) proteins. Center plots depict correlation links originating from only *GBA1*^*-/-*^ and *ASAH1*^*-/-*^. Only links with Pearson (r) smaller than -0.2 and larger than 0.7 are shown. Blue labels indicate genotypes belonging to Sphingolipidoses-risk genes. Bold labels are PD-risk genes.

### An Approach for Decoding Protein Complex-level Vulnerabilities

The experiments described above provide an atlas for evaluating the effects of LSD gene loss at an organelle level based on protein abundance. In order to extend this analysis to the level of individual PPIs and protein complexes, we developed an analysis pipeline that encompasses three major types of interaction data in a master PPI database: 1) well-validated protein complexes as are available in CORUM, Complex Portal (CP) or protein structure databases (PDB) (27-29), 2) high throughput interaction data, as represented in BioGrid and STRING databases (30, 31), and 3) predicted pairwise PPIs based on large-scale Alphafold Multimer or coevolutionary analyses (32, 33) (see **Fig. 4*A*, *SI Appendix*, Materials and Methods, Dataset S4**). Combining these resources with supporting structural evidence allowed us to generate a “baseline” neuronal cell line PPI network based on Tier 1 confidence scores (**Fig. 4*A***), ensuring that only high-quality predictions would be used for the downstream biological analysis. Over half (55.9%) of all edges in the neuronal baseline dataset were matched to structures in PDB, while nearly a quarter were also found in either CORUM or CP (***SI Appendix*, Fig. S4*A***). Of the ∼7100 PPIs found in the baseline database only 245 (∼3%) had no additional structural or biological validation beyond the very high protein structure prediction metrics. To assess the combinatorial overlap of supporting evidence in a numerical fashion, we developed a cumulative score capturing the structural and biological evidence for each PPI (***SI Appendix*, Fig. S4*B***,***C***). Not surprisingly, the highest counts per annotation were measured in mitochondrial, synaptic and nuclear annotations, thus scaling with both the number of proteins within a given annotation and the experimental evidence available in databases (***SI Appendix*, Fig. S4*D***,***E***,***F***). To normalize for the varying size of annotations we expressed the coverage of PPI edges in support databases as a fraction of the size of all edges found in the annotation. Compartments with extensive structural and biological characterization (e.g. mitochondria) displayed complete coverage across all sets, whereas structurally less-well defined compartments (e.g. ER) scored more poorly (***SI Appendix*, Fig. S4*D***, *bottom*). To expand this network further, we used the approach of Zhang et al (33) to first build 2571 and 1695 additional trimers and tetramers, respectively, from pairwise PPIs, with AF-Mprob >0.995, and >74% of these exhibiting structural support in the PDB (**Fig. 4*A-C*, *SI Appendix*, Fig. S4*G***,***H***,***I***). Fusion of high-confidence PPIs to at least 2 nodes of pre-existing trimers and tetramers seeds resulted in 408 complexes with a median support score of 4.06 across all databases (see ***SI Appendix*, Materials and Methods** for details of confidence limits). While 246 (60.3%) of these matched one or more high confidence complexes, 138 represent candidate new assemblies not found in existing structural databases but display high support metrics from structure prediction algorithms (**Fig. 4*D-F*, *SI Appendix*, Fig. S4*J***). In the identified complexes, between 45-60% of edges were matched to pre-existing structural database evidence, suggesting robust empirical support for these structures (**Fig.4*E,F***).

**Fig. 4.**
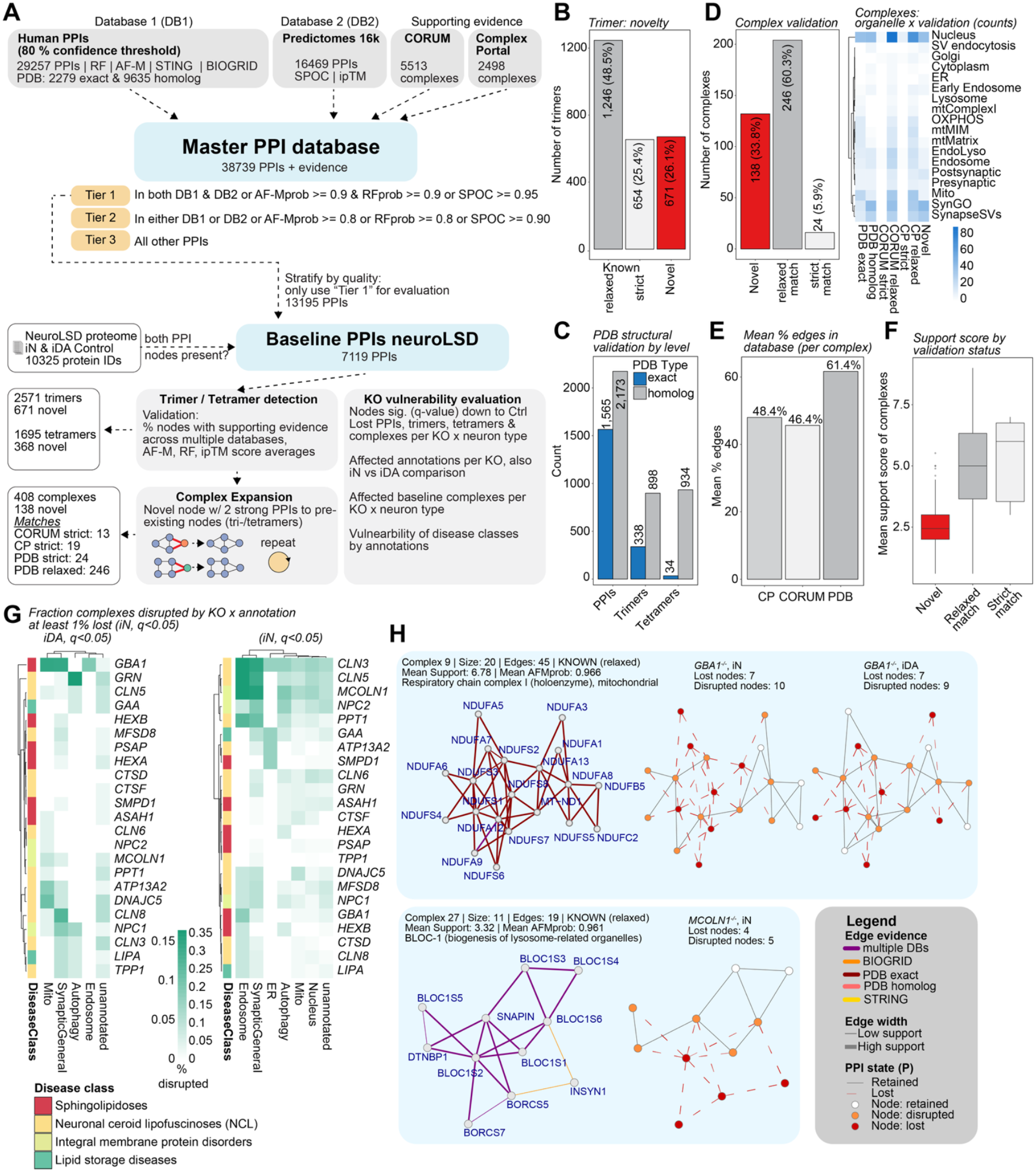
Protein-Protein interaction atlas of neuronal lysosomal dysfunction. ***(A)*** Schematic depiction of data evaluation pipeline for neuronal protein-protein interactions (PPIs) in LSD mutants. Only Tier 1 interactions were used for protein complex vulnerability analysis. ***(B)*** Bar graph of trimers found in neuronal baseline PPIs matched to pre-existing complex and structural databases. ***(C)*** Bar graph showing number of PPIs, trimers and tetramers (from left to right) with matching structural validation in PDB database. ***(D)*** Like panel **B**, but for complexes detected in neuronal baseline PPI dataset. Right panel displays a heatmap of the counts of validated complexes across organelles. ***(E)*** Bar graph of mean percent of PPI edges found in ComplexPortal (CP), CORUM and PDB. ***(F)*** Boxplot of mean support score of complexes belonging to novel, relaxed match and strict match classes. ***(G)*** Heatmap of fraction of PPIs (q < 0.05) disrupted by organelle across LSD KOs in iN (left) and iDA (right). ***(H)*** Examples representation of protein complex vulnerability. (top) Mitochondrial complex I is depicted as example with loss or disruption of several nodes in *GBA1*^*-/-*^ iN (middle) or *GBA1*^*-/-*^ iDA (right) compared with Control cells (left). (bottom left) BLOC-1 complex as an example of lost or disrupted interactions in *MCOLN1*^-/-^ iN cells. The legend summarizes color codes for edge evidence and PPI state.

Having developed this protein interaction database, we first identified individual pairwise PPIs where the abundance of one or both proteins was significantly reduced (q < 0.05) compared with controls across LSD genotypes (***SI Appendix*, Fig. S5*A***). The majority of pairwise interactions were unaffected by LSD gene mutation, although genotypes displaying the largest number of proteome alterations (e.g. CLN3, CLN5, CLN6 and MCOLN1) displayed the highest fractional PPI reduction, as expected. Reduced pairwise interactions were enriched in annotations for specific subcellular compartments in a genotype-dependent and in some cases, disease class-dependent manner in both iN and iDA cells, suggesting selective vulnerability (**Fig. 4*G, SI Appendix*, Fig. S5*B***). To map proteome alterations at the protein complex level (i.e. beyond pairwise interactions), we assigned PPI nodes within complexes as “disrupted” when the abundance of at least one interaction (edge) was significantly reduced and as “lost” when *all* detected interactions with that node displayed reduced abundance (**Fig. 4*A**, SI Appendix*, Fig. S5*C* and Dataset S4**) The number of lost or disrupted nodes varied across LSD genotypes and was generally more prevalent in iN cells (***SI Appendix*, Fig. S5*C* and Dataset S4**). Additionally, individual LSD genotypes displayed enrichment of specific organelles for disrupted or lost protein complexes, including trimers built using the method of Zhang et al (33) (**Fig. 4*G**, SI Appendix*, Fig. S5*D* and Dataset S4**).

Examples of disrupted complexes are shown in **Fig. 4*H* *and SI Appendix*, Fig. S5*E***. One robust case is Complex I from the mitochondrial electron transport chain, which displayed 7 or 10 lost or disrupted nodes, respectively, in *GBA1*^-/-^ iN or iDA cells, consistent with enrichment for OXPHOS localization terms being affected at the organelle level (**Fig. 2*D***). Similarly, the BLOC-1 (Biogenesis of lysosome-related organelles) complex known to regulate lysosome position in cells (34) displayed loss or disruption of 4 or 5 nodes in *MCOLN1*^-/-^ iN cells. Additional examples of altered complexes include: 1) the core Commander Complex involved in sorting of proteins into recycling vesicles (35) displays 9 disrupted nodes in *NPC1*^-/-^ iN cells, 2) glutamate receptor assemblies (36) in *GBA1*^-/-^ and *MCOLN1*^-/-^ cells, 3) CAMK2 in *GBA1*^-/-^, *ASAH1*^-/-^ and *MCOLN1*^-/-^ cells, 4) the GATOR complex involved in amino acid sensing (37) in *ASAH1*^-/-^ cells, 5) the electron transport chain complex cytochrome C oxidase in *ASAH1*^-/-^ cells, and 6) the RAB7 guanine nucleotide exchange factor CCZ1-MON1-RMC1 on lysosomes (38-40) in *ASAH1*^-/-^ cells (***SI Appendix* Fig. S5*E***). The entire collection of protein complexes with alterations in specific subunits is available in **Dataset S4**. Taken together, these examples demonstrate how the interaction analysis pipeline allows interrogation of altered proteomes beyond the level of organelles in a systematic manner.

### Synaptic Function in *ASAH1*^-/-^ iDA cells

As described above, we detected alterations in synaptic protein abundance in *ASAH1*^-/-^ iNeurons, which led us to examine synaptic structure and function in these cells. We initially performed thin-section electron microscopy on day 50 Control and *ASAH1*^-/-^ iDA and iN cells (**Fig. 5*A, SI Appendix*, Fig. S6*A***). In Control iDA cells, we observed pre-synaptic structures containing numerous synaptic vesicles with uniform vesicle sizes (**Fig. 5*A***). In contrast, apparent synaptic structures in *ASAH1*^-/-^ iDA cells appeared more disorganized with often fewer synaptic vesicles and with less uniform size distributions (**Fig. 5*A***). Similar findings were made in day 50 iN cells (***SI Appendix*, Fig. S6*A***). Immunostaining of day 38 iDA cultures with the pre-synaptic markers SYP (synaptophysin) and BSN (Bassoon) revealed largely uniform pre-synaptic with the majority of SYP puncta co-staining with Bassoon in Control cells (**Fig. 5*B-D**, SI Appendix*, Fig. S6*B***). In contrast, *ASAH1*^-/-^iDA cells frequently displayed enlarged α-tubulin positive neuronal structures that contained α-SYP puncta that was more frequently not co-stained with α-Bassoon (**Fig. 5*B-D**, SI Appendix*, Fig. S6*A***), suggesting defects in pre-synaptic compartment organization.

**Fig. 5.**
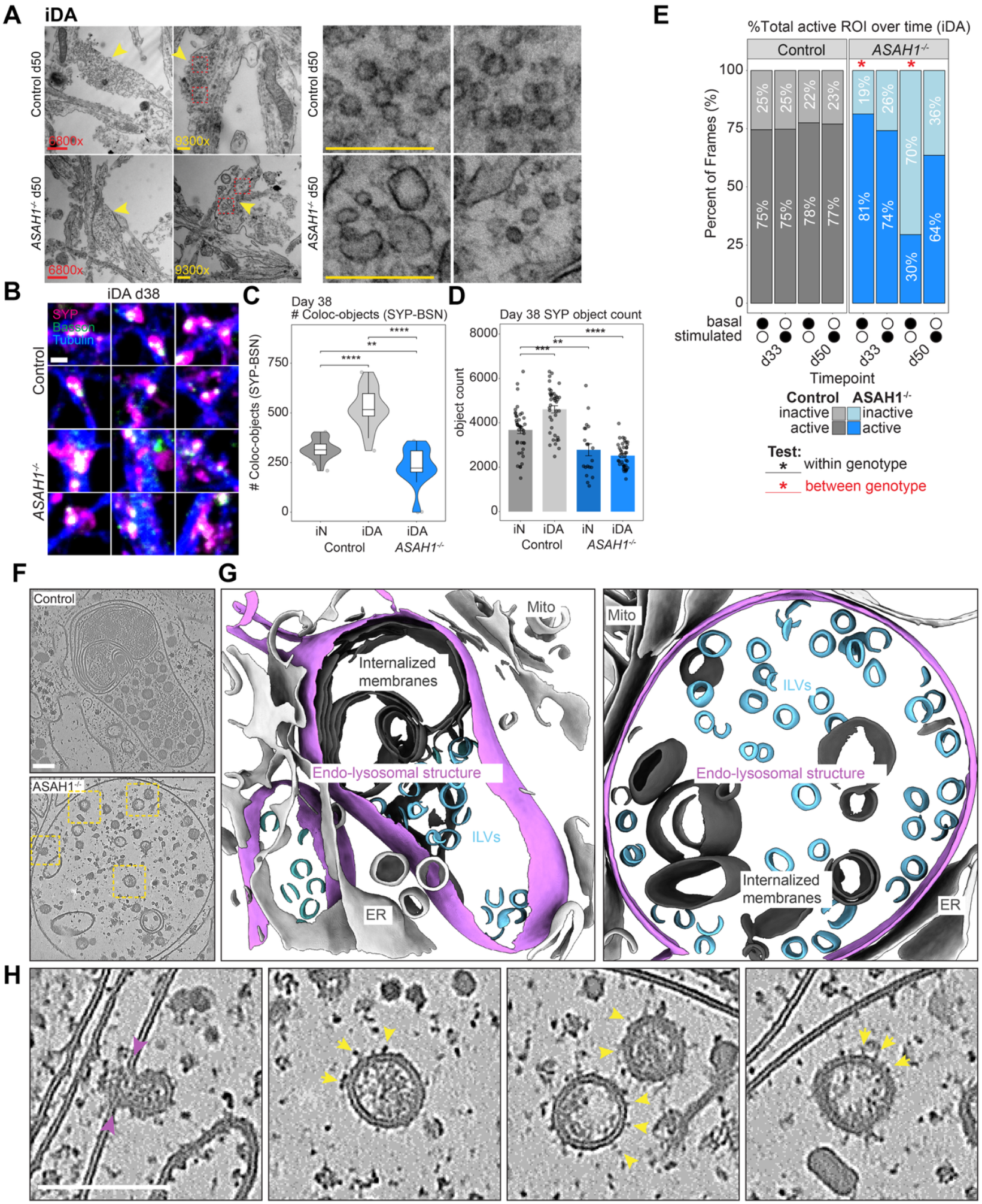
Synaptic morphology defects correlate with reduced basal Ca^2+^ firing activity in *ASAH1*^-/-^ iDA cells. **(*A*)** Representative EM images depicting synapses and neuronal ultrastructure of Control and *ASAH1*^*-/-*^ iDA neurons at day 50 of *in vitro* differentiation. Scale bars are indicated. Pre-synaptic structures with synaptic vesicles are indicated by yellow arrowheads. Panels on the right display enlarged vesicular structures within *ASAH1*^-/-^ synapses. **(*B*)**, Confocal images of Control and *ASAH1*^*-/-*^ iN and iDA cells at day 38 of *in vitro* differentiation, labeled for SYP, Bassoon and Tubulin. Scale bar: 20 µm and 2 µm. Images depicting a wider field view are provided in **SI Appendix, Fig. S6*B*. (*C*)** Number of colocalized SYP-Bassoon object pairs in Control iN and iDA and iDA *ASAH1*^*-/-*^ neurons, based on confocal images in panel ***B*** (n(Control iN): 20 stacks; n(Control iDA): 19 stacks; n(*ASAH1*^*-/-*^ iDA): 19 stacks). iN: **P(0.001); iDA: ****P< 0.0001 ; iDA Ctrl vs *ASAH1*^*-/-*^: ****P< 0.0001. **(*D*)** Quantification of SYP object counts (from left to right), based on confocal images in panel ***B***. (n(Control iN): 40 stacks; n(Control iDA): 40 stacks; n(*ASAH1*^*-/-*^ iN): 20 stacks; n(*ASAH1*^*-/-*^ iDA): 40 stacks). Object count: Control iN vs iDA: ***P(0.0002), iN Control vs *ASAH1*^*-/-*^: **P(0.003), iDA Control vs *ASAH1*^*-/-*^: ****P<0.0001. **(*E*)** Stacked bar graph for Control (left) and *ASAH1*^*-/-*^ (right) iDA cell Ca^2+^ firing depicting the cumulative % of active ROIs over the time course. d33 Control vs *ASAH1*^*-/-*^: *P(0.02); d50 Control vs *ASAH1*^*-/-*^: *P(0.04). Data from day 33 and 50 of *in vitro* differentiation with or without stimulation by KCl. **(*F*)** Example tomogram of Control and *ASAH1*^*-/-*^ endo-lysosomal structures. Scale bar 100 nm. **(*G*)** 3D-renderings of tomogram segmentations from panel ***F*. (*H*)** Higher magnification of the white boxes in panel ***F*** showing inward budding membrane events (purple arrowheads) and ILVs studded with glycosylated proteins (yellow arrowheads). Scale bar 100 nm.

To examine the functional impact of *ASAH1* deficiency, we performed Fluo-4-based Ca^2+^ imaging of Control and *ASAH1*^-/-^ iDA and iN cells over increasing time periods of differentiation using a robust evaluation machine-learning-based pipeline (41, 42) (***SI Appendix*, Fig. S6*C-E***) (see Materials and Methods). As expected, stimulation of Control cells with KCL resulted in an increase in relative firing frequency, which could be reduced by pre-treatment with CQNX/D-AP5 (***SI Appendix*, Fig. S6*D*)**. We found that neuronal firing, as measured by either the percent of total active regions of interest (ROI) over time or the mean percentage of active versus non-active frames, were significantly reduced in day 50 iDA *ASAH1*^*-/-*^ vs Control cells, but this reduced activity could be partially rescued, albeit not to Control-levels, by the addition of KCl (**Fig. 5*E**, SI Appendix*, Fig. S6*E***). In contrast, firing rates were only minorly affected in iN cells over the selected timepoints (***SI Appendix*, Fig. S6*F***). Thus, alterations in the synaptic proteome in day 50 iDA cells correlates with reduced firing rates.

### Visualization of *ASAH1*^-/-^ endolysosomal compartments by cryo-ET

ASAH1 plays a central role in glycosphingolipid catabolism within the lysosome through the conversion of ceramide to sphingosine and fatty acids and functions downstream of GBA1 and upstream of SMPD1 (***SI Appendix*, Fig. S7*A***). The finding that *ASAH1*^-/-^ iDA and iN cells display altered lysosomal proteome abundance (**Fig. 3*A,B***) prompted a biochemical and ultrastructural analysis of ASAH1-deficient lysosomes in the experimentally tractable HeLa cells system. Initially, we performed LysoIP from Control and *ASAH1*^-/-^ HeLa cells containing an endogenously tagged TMEM192^HA^ locus (4) along with parental HeLa cells as a negative control. Proteomic analysis of isolated lysosomes displayed the expected increase in endolysosomal proteins, which were absent in cells lacking the LysoIP tag (***SI Appendix*, Fig. S7*B***,***C* and Dataset S5**), with a subset of lumenal hydrolytic proteins displaying increased abundance (***SI Appendix*, Fig. S7*D***). As expected, lipidomics revealed a ∼3-fold increase in ceramide abundance in lysosomes from *ASAH1*^-/-^ cells, consistent with *ASAH1* deficiency (***SI Appendix*, Fig. S7*E* and Dataset S6**). Interestingly, several BMP species were increased in abundance in lysosomes in the absence of *ASAH1*, which may underlie the modest (∼2-fold or less) reduction in glycosphingolipids (GSL) GM1, GM2, GM3 in lysosomes (7) (***SI Appendix*, Fig. S7*F***). Taken together, this analysis validates the *ASAH1*^-/-^ cell line and is consistent with its role as an acid ceramidase.

We next examined endolysosomal structures *in situ* in *ASAH1*^-/-^ HeLa cells using cryo-focused ion beam (cryo-FIB) milling paired with cryo-ET (**Fig. *5F-H***). Endolysosomal structures in Control cells have typical ultrastructural features of auto-/endolysosomes, including frequent single or double membrane structures within the limiting membrane and dense material indicative of degradative cargo (**Fig. 5*F,G*)**. Endolysosomal structures in *ASAH1*^-/-^ cells appeared swollen compared to those in Control cells, which showed dense packing of cellular material (**Fig. 5*F,G*)**. Notably, membranous material was largely absent in endolysosomal structures in *ASAH*^*-/-*^ cells. In one tomogram of *ASAH1*^-/-^ cells, we observed an inward-budding vesicle within an endolysosome, along with numerous intraluminal vesicles (ILVs) that appeared to be studded with glycosylated proteins (**Fig 5*F-H*** and additional examples in ***SI Appendix*, Fig. S8*A***,***B***). ILVs are thought to be the sites of breakdown of glycosphingolipids (5, 8). Segmentation of tomograms highlights the difference in size and membrane structure packing within the endolysosomes of Control or *ASAH1*^-/-^ cells, and reveals the close proximity of ER to the limiting membrane (**Fig. 5*G**, SI Appendix*, Movie S1, S2**). Taken together, these results suggest endolysosomal dysfunction in the context of *ASAH1* deficiency.

## Discussion

The LSD toolkit reported here provides a means by which to examine consequences of individual LSD gene loss during changes in cell states while also allowing comparisons across various classes of storage disorders. Through extensive analysis of total proteome data for both iN and iDA cells, we provided a landscape of alterations reflecting the activities of individual LSD genes. To our knowledge, this is the first comparison between iN and iDA cells in culture at this level of proteome depth, and the first comparison across divergent LSD mutants. Additionally, we developed an informatic framework to globally map alterations in specific protein complexes based on alterations in protein abundance. This PPI approach allows us to go beyond protein abundance changes at the organelle level to map alterations in the context of protein complexes, including potentially new interactions based on proteome-wide computational structure prediction methods. Although deployed here on neuronal proteomes across LSD mutations, the approach should be adaptable for mapping protein complex alterations across virtually any large-scale data types, including CRISPR, Perturb-seq or degrader screens. Additionally, our analysis focused on Tier 1 PPIs, which represents the most stringent evidence for validated interactions. Further analysis, including Tier 2 complexes (**Fig. 4*A***), may reveal additional complexes of interest whose abundance may be altered in specific genetic contexts. Our ongoing studies seek to expand the toolkit to ultimately encompass all genes associated with LSDs. To facilitate accessibility of the proteomic data, we provide an interactive data viewer at https://wren.hms.harvard.edu/ProteomeLSDNeuron/.

Our analysis focused on genes linked with sphingolipidoses, in part due to potential links with PD risk (18). We found that *GBA1* and *ASAH1* mutants sometimes displayed distinct proteome alterations compared with other sphingolipidoses genes. In particular, we observed alternations in OXPHOS and pre-synaptic proteins in *GBA1* and *ASAH1* mutant iDA cells. The selectivity observed for alterations in the mitochondrial OXPHOS system in iDA cells is interesting in light of the known linkage between OXPHOS pathways and PD (43, 44). Conditional deletion of murine mtComplex I subunit *Ndufs4* results in a deficit in nigrostriatal axons, and is associated with a PD-like loss of motor function, suggesting that loss of mtComplex I component of OXPHOS is sufficient to produce motor dysfunction (44). Likewise, the mtComplex I inhibitors MPTP and rotenone promote a Parkinson’s-like phenotype in mice (45, 46). Similarly, synaptic dysfunction has also been linked with PD, including synaptotagmin-11 (*SYT11*) and synaptojanin (*SYNJ1*) variants that have been associated with familial PD (47, 48). It is conceivable that alterations in OXPHOS and pre-synaptic protein pools observed in *ASAH1*^-/-^ and *GBA1*^-/-^ iDA cells could represent vulnerabilities that increase the risk of PD in patients with reduced enzymatic activity for these key sphingolipid catabolism enzymes. Loss of *ASAH1* results in increased levels of ceramide, as confirmed here by lipidomics in HeLa cells. Ceramides are known to affect the electron transport chain in mitochondria (49), suggesting additional potential mechanisms independent of alterations in OXPHOS protein abundance. Interestingly, we also observed increased levels of BMP species in *ASAH1*^-/-^ HeLa cells. We speculate that increased levels of BMP may increase the efficiency of catabolism of sphingolipids in the context of *ASAH1* mutants, as indicated by reduced levels of GM2 and GM3 species. Future studies are required to understand the relative contributions of alterations in synaptic and mitochondrial protein abundance for calcium firing rate effects seen in *ASAH1*^-/-^ iDA cells.

We note several limitations of the current study: 1) Our analysis employed deletion mutants rather than patient mutants, and therefore the phenotypes observed may be more severe than with risk alleles observed in patients; 2) Our analysis was performed in monocultures, and further studies will be required to understand if co-culture with astrocytes or glial cells alters observed phenotypes; 3) All of our analyses of vulnerabilities for protein interactions employed Tier 1 interactions and therefore represent the most stringent large-scale data available, but directed quantitative studies would be necessary to validate loss of particular protein complexes, which might also be cell type dependent; 4) We employed HeLa cells for analysis of *ASAH1*^-/-^ lysosomes by cryo-ET given the challenges associated with analysis of neurons by this technique, and alterations in lysosomal structure will need to be validated further in more physiologically relevant cells; and 5) It is possible that certain alterations within specific compartments (e.g. lysosomes) are missed by the whole cell analysis approach used here but future studies using organelle enrichment methods, including the use of the Lyso-IP tag incorporated into the ES cell system used here, would provide an approach to examine both proteome and lipidome alterations within lysosomes in order to better understand the effects of removal of specific LSD genes.

## Materials and Methods

Detailed procedures related to cell culture, gene editing, biochemical procedures, MS, microscopy, and electron tomography are described in *SI Appendix, Materials and Methods*. Statistical analyses were performed using MSstats and R. All error bars represent SEM, and statistical significance was determined by *t* tests, as specified in the corresponding figure legends.

## Data, Materials, and Software Availability

Proteomic data (.RAW files) for nDIA of iN/iDA day 30, 50 and 70 iNeurons is deposited in the MASSIVE Database (https://massive.ucsd.edu/ProteoSAFe/static/massive.jsp): MSV000099237 (reviewer login: MSV000099237_reviewer; password: lsd). HeLa LysoIP TMT data was deposited into ProteomeXchange (50): PXD067219. Lipidomics data is deposited on Metabolomics Workbench (51): Study ID ST004217; http://dx.doi.org/10.21228/M8Z556. Raw Cryo-ET tomograms have been deposited at EMDB under accession numbers EMD-55210 (Control) and EMD-55211 (*ASAH1*^-/-^). A Key Resource Table containing reagents and materials, source data, segmentation models and datasets associated with this publication are available from Zenodo.org: 10.5281/zenodo.17296003. The Key Resource Table also contains unique Cellosaurus (https://www.cellosaurus.org/) identifiers for all cell lines reported here, which are available upon request with requisite Material Transfer Agreements from WiCell. Scripts affiliated with this work is available on Github under the repository https://github.com/sauerkrausi/neuroLSD.

Data viewer can be found at: https://wren.hms.harvard.edu/ProteomeLSDNeuron/.

## Acknowledgments

We thank members of the Harper lab for feedback. We thank the Central Electron Microscopy Facility at the Max Planck Institute of Biophysics for support. This work was funded by the Warren Alpert Foundation (J.W.H.), the Bluefield Project (J.W.H., R.V.F, T.C.W.), NIH R01NS110395 (J.W.H.), NIH R01 GM132129 (J.A.P.), the Max Planck Society (F.W.), a Fred and Joan Goldberg Post-doctoral Fellowship (F.K.). D.L. was supported by a Boehringer Ingelheim Fonds PhD fellowship. This research was also funded by Aligning Science Across Parkinson’s [ASAP-024268 and 025160 to J.W.H. and F.W.] through the Michael J. Fox Foundation for Parkinson’s Research (MJFF). For the purpose of open access, the author has applied a CC BY public copyright license to all Author Accepted Manuscripts arising from this submission. We acknowledge the Core for Imaging Technology and Education (CITE, Harvard Medical School) for imaging assistance. T.C.W. is a Howard Hughes Medical Institute investigator.

