## Supplementary material for "A human lysosomal storage disorder toolkit for decoding proteome landscapes in cortical and dopaminergic-like induced neurons": SI Appendix

^Current address: Department of Pathology, University of British Columbia, Vancouver, BC, Canada.

**This PDF file includes:**

Materials and Methods
Figures S1 to S8
Legends for Movies S1 and S2
Legends for Datasets S1 to S6
SI References

**Other supporting materials for this manuscript include the following:**

Movies S1 to S2
Datasets S1 to S6

### Supporting Information Text

### MATERIALS and METHODS

Details of key resources used in this study are provided in a Key Resource Table available on Zenodo: 10.5281/zenodo.17296003.

#### Reagents

The following chemicals and reagents were used: 100x21mm Dish, Nunclon Delta (Thermo Fisher Scientific, 172931); 12 Well glass bottom plate with high performance #1.5 cover glass (Cellvis, P12-1.5H-N); 150 mm plates, 15 cm (MidSci, TP93150); 16% Paraformaldehyde, Electron-Microscopy Grade (Electron Microscopy Science, 15710); 2-Chloroacetamide (Sigma-Aldrich, C0267); 24 Well glass bottom plate with high performance #1.5 cover glass (Cellvis, P24-1.5H-N); 6 Well glass bottom plate with high performance #1.5 cover glass (Cellvis, P06-1.5H-N); 96 Well glass bottom plate with high performance #1.5 cover glass (Cellvis, P96-1.5H-N); Accutase (StemCell Technologies, 7920); Acetonitrile, Optima LC/MS Grade (Thermo Fisher Scientific, A955-4); Adenosine 5′ triphosphate, disodium, trihydrate (Thermo Fisher Scientific, 10326943); Ammonium formate CHROMASOLV LC-MS Ultra (Honeywell, 14266 Fluka); Anti-Flag M2 magnetic beads (Sigma Millipore, M8823; RRID: AB_2637089); B27 (Thermo Fisher Scientific, 17504001); Brain-derived neurotrophic factor, BDNF (PeproTech, 450-02); CloneR (StemCell Technologies, 5889); CNQX (Cayman, 14618); cOmplete, EDTA-free Protease Inhibitor Cocktail (Millipore Sigma-Aldrich, 11873580001); Corning Matrigel Matrix (Corning, 354230); Corning 24 mm Transwell with 3.0 µm Pore Polycarbonate Membrane Insert, Sterile (Corning, 3414); Cover Glasses (VWR, 16004-308); Cultrex 3D Culture Matrix Laminin I (R&D Systems, 3446-005-01); D-AP5 (MedChemExpress, HY-100714A); DAPI (Thermo Fisher Scientific, D1306); Dextran, Alexa Fluor 647; 10 000 MW, Anionic, Fixable (Thermo Fisher Scientific, D22914); Dithiothreitol, DTT (Gold Biotechnology, DTT25); DMEM, High Glucose, Pyruvate (Thermo Fisher Scientific, 11995-073); DMEM/F12 (Thermo Fisher Scientific, 11330057); DNAse I (Thermo Fisher Scientific, EN0521); dNTPs (New England Biolabs, N0447L); Dounce homogenizer (DWK Life Sciences, 885302-0002); Doxycycline (Sigma-Aldrich, D9891); EPPS (Sigma-Aldrich, E9502); Fetal bovine serum (Cytiva, SH30910.03); FGF3 (in-house, N/A); Fluo-4, AM, cell permeant (Thermo Fisher Scientific, F14201); FluoroBrite DMEM (Thermo Fisher Scientific, A1896701); Formic Acid (Sigma-Aldrich, 94318); Geltrex LDEV-Free Reduced Growth Factor Basement Membrane Matrix (Thermo Fisher Scientific, A1413202); GlutaMAX (Thermo Fisher Scientific, 35050061); High-pH fractionation kit (Thermo Fisher Scientific, 84868); Hoechst33342 (Thermo Fisher Scientific, H1399); Holo-transferrin, human (Sigma-Aldrich, T0665); Human EGF Recombinant Protein (Cell Signaling Technology, 72528S); Human GDNF Recombinant Protein (PeproTech, 450-10-50UG); Human insulin (Sigma-Aldrich, I9278-5ML); Hydroxylamine solution (Sigma-Aldrich, 438227); Hygromycin B (Thermo Fisher Scientific, 10687010); Immobilon-P Membrane, PVDF, 0.45 µm (EMD Millipore, IPVH00010); Isopropanol, Optima LC/MS Grade (Thermo Fisher Scientific, A461-4); Lipofectamine LTX (Thermo Fisher Scientific, 15338100); Lys-C (Wako Chemicals, 129-02541); LysoTracker Red DND-99 (Thermo Fisher Scientific, L7528); MEM NEAA (Thermo Fisher Scientific, 11140050); N-2 Supplement (Thermo Fisher Scientific, 17502048); n-Butanol (Thermo Fisher Scientific, A383SK-4); NEAA (Life Technologies, 11140050); NEBNext Ultra II Q5 Master Mix (New England Biolabs, M0544L); Neurobasal (Thermo Fisher Scientific, 21103049); Neurotrophin-3, NT3 (PeproTech, 450-03); Nunc 12-well plate (Thermo Fisher Scientific, 150628); Nunc Cell-Culture Treated 6-well (Thermo Fisher Scientific, 140685); NuPAGE Novex 4-12% Bis-Tris Midi Protein Gels, 20 well (Thermo Fisher Scientific, WG1402BOX); NuPAGE Novex 4-12% Bis-Tris Midi Protein Gels, 26 well (Thermo Fisher Scientific, WG1403BOX); NuPAGE LDS Sample Buffer, 4X (Thermo Fisher Scientific, NP0008); Oligo dT20 primers (Invitrogen, 79654); OptiMEM I Reduced Serum Media (Thermo Fisher Scientific, 31985062); PBS (Corning, 21-031-CV); Penicillin-Streptomycin, 10 000 U/mL (Thermo Fisher Scientific, 15140163); PhosSTOP Phosphatase Inhibitor Cocktail (Roche, 4906845001); Pierce anti-HA magnetic beads (Thermo Fisher Scientific, 88837); Poly-L-ornithine hydrobromide (Sigma-Aldrich, P3655-50MG); Polyethylenimine (Polysciences, 23966); Precision Plus Protein Kaleidoscope Prestained Protein Standards (Bio-Rad, 1610395); Puromycin (Gold Biotechnology, P-600-500); Recombinant Human Sonic Hedgehog, Shh (PeproTech, 100-45); Recombinant SpCas9 (in-house, N/A); Revert 700 Total Protein Stain Kit (LI-COR, 926-11016); SDS (Bio-Rad, 1610302); Sep-Pak C18 cartridge (Waters, WAT054955); Sep-Pak C18 1cc Vac Cartridge, 50 mg (Waters, WAT054960); Sep-Pak tC18 96-well Plate, 25 mg Sorbent per Well (Waters, 186002319); SeraSil-Mag silica-coated superparamagnetic beads, 700 nm (Cytiva, 29357374); Silicon Dioxide film (Quantifoil,

50-192-7684); Silicon Dioxide R1/4 film (Quantifoil, 50-192-7684); SiR-Lysosome Kit (Cytoskeleton / Spirochrome, CY-SC012); SiR-Tubulin Kit (Cytoskeleton / Spirochrome, CY-SC002); Sodium bicarbonate (Sigma-Aldrich, S5761-500G); Sodium Pyruvate (Invitrogen, 11360070); Sodium selenite (Sigma-Aldrich, S5261-10G); SOLA HRP SPE Cartridge, 10 mg (Thermo Fisher Scientific, 60109-001); SOLA HRP SPE 30mg/2mL 96-well plate (Thermo Fisher Scientific, 60509-001); SPLAH Lipidomix Mass Spec Standard (Avanti, 330707-1EA); STEMdiff Midbrain Neuron Differentiation Kit (StemCell Technologies, 100-0038); STEMdiff Midbrain Neuron Maturation Kit (StemCell Technologies, 100-0041); StrataX 10 mg 96-well Plate (Phenomenex, 8E-S100-AGB); TCEP (Gold Biotechnology, TCEP2); TGF-beta (PeproTech, 100-21C); TMTpro 18plex Label Reagent (Thermo Fisher Scientific, A52045); Tris 1 M pH 8.0 RNase-free (Thermo Fisher Scientific, AM9855G); Tris(2-carboxyethyl)phosphine, TCEP (Gold Biotechnology, 51805-45-9); Trypsin (Promega, V511C); Trypsin-EDTA (Sigma-Aldrich, T4049-100ML); UltraPure 0.5 M EDTA pH 8.0 (Thermo Fisher Scientific, 15575020); Urea (Sigma-Aldrich, U5378); Uridine (Sigma-Aldrich, U3003); Vectashield (Vector Laboratories, H-1000-10); Water, Optima LC/MS Grade (Thermo Fisher Scientific, W64); Y-27632 Dihydrochloride ROCK inhibitor (PeproTech, 1293823); μ-Slide 8 Well Glass Bottom #1.5H Coverslip (ibidi, 80807); SpCas9(1) and AsCas12a/AsCpf1(2) were from a previous study (3).

The following antibodies were used: Anti-Actin (Sigma-Aldrich, A2228; RRID:AB_476697); Anti-Bassoon monoclonal antibody, SAP7F407 (Enzo, ADI-VAM-PS003-D); Anti-EEA1, C45B10 Rabbit mAb (Cell Signaling Technology, 3288S); Anti-Flag antibody (Sigma-Aldrich, F1804); Anti-HA antibody (Cell Signaling Technology, 3724); Anti-HA High Affinity, rat IgG1 (Roche, 11867423001; RRID:AB_390918); Anti-LAMP1, D2D11 XP Rabbit (Cell Signaling Technology, 9091S; RRID:AB_2687579); Anti-LAMP1, D4O1S Mouse (Cell Signaling Technology, 15665; RRID:AB_2798750); Anti-MAP2, D7H11 XP Rabbit mAb (Cell Signaling Technology, 4542S); Anti-mouse IgG HRP conjugate (Bio-Rad, 170-6516; RRID:AB_11125547); Anti-Neurofilament heavy polypeptide antibody (Abcam, ab4680); Anti-rabbit IgG HRP conjugate (Bio-Rad, 170-6515; RRID:AB_11125142); Anti-Synapsin I (SYN1) antibody [EPR23531-50], recombinant (Abcam, ab254349); Anti-Synaptophysin antibody [YE269] (Abcam, ab32127); Anti-Tubulin beta III antibody, chicken (Abcam, ab41489; RRID:AB_727049); Anti-Tyrosine Hydroxylase Antibody, clone LNC1 (Sigma-Aldrich, MAB318); Donkey anti-Rat IgG (H+L) Highly Cross-Adsorbed Secondary Antibody, Alexa Fluor Plus 405 (Thermo Fisher Scientific, A48268; RRID:AB_2890549); Goat anti-Chicken IgY (H+L) Secondary Antibody, Alexa Fluor 488 (Thermo Fisher Scientific, A-11039; RRID:AB_2534096); Goat anti-Chicken IgY (H+L) Secondary Antibody, Alexa Fluor 647 (Thermo Fisher Scientific, A-21449; RRID:AB_2535866); Goat anti-Mouse IgG (H+L) Cross-Adsorbed Secondary Antibody, Alexa Fluor 488 (Thermo Fisher Scientific, A-11001; RRID:AB_2534069); Goat anti-Mouse IgG (H+L) Cross-Adsorbed Secondary Antibody, Alexa Fluor 647 (Thermo Fisher Scientific, A-21235; RRID:AB_2535804); Goat anti-Rabbit IgG (H+L) Cross-Adsorbed Secondary Antibody, Alexa Fluor 568 (Thermo Fisher Scientific, A-11011; RRID:AB_143157); Goat anti-Rabbit IgG (H+L) Cross-Adsorbed Secondary Antibody, Alexa Fluor 647 (Thermo Fisher Scientific, A27040; RRID:AB_2536101); Goat anti-Rabbit IgG (H+L) Highly Cross-Adsorbed Secondary Antibody, Alexa Fluor Plus 405 (Thermo Fisher Scientific, A48254; RRID:AB_2890548); Goat anti-Rabbit IgG (H+L) Highly Cross-Adsorbed Secondary Antibody, Alexa Fluor 488 (Thermo Fisher Scientific, A-11034; RRID:AB_2576217); Goat anti-Rat IgG (H+L) Cross-Adsorbed Secondary Antibody, Alexa Fluor 555 (Thermo Fisher Scientific, A-21434; RRID:AB_2535855); Goat anti-Rat IgG (H+L) Cross-Adsorbed Secondary Antibody, Alexa Fluor 647 (Thermo Fisher Scientific, A-21247; RRID:AB_141778). All primary antibodies were used 1:500 for IFA and 1:1000 for WB (if not otherwise stated); secondary antibodies were used 1:200 for IFA and 1:5000 for WB.

**Cell culture of HeLa cell lines**

HeLa TMEM192-3xHA cells (referred to as $HeLa^{TMEM192-HA}$) (3) were maintained in Dulbecco's modified Eagle's medium (DMEM), supplemented with 10% vol/vol fetal bovine serum (FBS), 5% vol/vol penicillin-streptomycin (P/S), 5% vol/vol GlutaMAX and 5% vol/vol non-essential amino acids (NEAA) at 37°C, 5% $O_2$, as described in the protocol: dx.doi.org/10.17504/protocols.io.kxygx31oog8j/v1. Unless otherwise noted, we refer to independently grown and handled cultures as biological replicates to distinguish from assays performed on identical samples (i.e. technical replicates).

**Stem-cell culture and neuronal differentiation**

Detailed methods for maintenance and differentiation of stem cells can be found at: dx.doi.org/10.17504/protocols.io.br85m9y6, dx.doi.org/10.17504/protocols.io.br9cm92w, dx.doi.org/10.17504/protocols.io.br9em93e, dx.doi.org/10.17504/protocols.io.bsacnaaw. Briefly, human

ES cells (H9, WiCell Institute) were cultured in E8 medium (4, 5) on Geltrex-coated tissue culture plates with daily medium change. Cells were passaged every 4-5 days with 0.5 mM EDTA in 1× DPBS (Thermo Fisher Scientific).

For conversion of human stem cells to iNeurons, cells were expanded and plated at $2x10^4/cm^2$ on Geltrex-coated tissue plates in DMEM/F12 supplemented with 1x N2, 1x NEAA (Thermo Fisher Scientific), human Brain-derived neurotrophic factor (BDNF, 10 ng/ml, PeproTech), human Neurotrophin-3 (NT-3, 10 ng/ml, PeproTech), mouse laminin (0.2 µg/ml, Cultrex), Y-27632 (10 µM, PeproTech) and Doxycycline (2 µg/ml, Alfa Aesar) on Day 0. On Day 1, Y-27632 was withdrawn. On Day 2, medium was replaced with Neurobasal medium supplemented with 1x B27 and 1x Glutamax (Thermo Fisher Scientific) containing BDNF, NT-3 and 1 µg/ml Doxycycline. Starting on Day 4, half of the medium was replaced every other day thereafter. On Day 7, the cells were treated with 0.5µM EDTA + PBS and plated on Geltrex-coated tissue plates. Following day 12 of differentiation 50% ND2 media was changed every other day. Cells were replated at day 16 into triple-coated dishes (Geltrex + poly-L-orthinine + laminin). After day 26, 50% of media was changed every 4 days.

For human ES cell conversion to induced dopaminergic neurons (iDA neurons), were differentiated as described(6). Cells were seeded on Geltrex-coated plates at DIV-2 and expanded in standard culture medium. On Day 0 (DIV 0), medium was replaced with DMEM/F12-based D0/1 induction medium supplemented containing N2, B27, NEAA, BDNF, GDNF, mouse laminin, and doxycycline (2 µg/ml) to induce NGN2 expression. On Day 1, the D0/1 induction medium (+dox) was refreshed. On Day 2 (DIV 2), cells were washed with PBS and switched to iDA differentiation medium supplemented with 2 µM Ara-C to inhibit proliferating cells. Media was changed daily from DIV 2 to DIV 4. From DIV 5 to DIV 8, half of the medium was replaced daily with fresh iDA differentiation medium. On Day 9, cells were transitioned to midbrain maturation medium containing doxycycline. From DIV 10 onward, half of the maturation medium was replaced every other day until terminal differentiation (e.g., DIV 26). Cells were replated at day 16 into triple-coated dishes (Geltrex + poly-L-ornithine+ laminin). After day 26, 50% of media was changed every 4 days.

**Gene-Editing**

Our protocol for gene editing of LSDs can be found at: dx.doi.org/10.17504/protocols.io.dm6gpmdwpgzp/v1. Generation of LSD mutants in $H9^{NGN2;TMEM192\text{-}HA}$ cells(3, 7) was facilitated using CRISPR/Cas9 with target sites determined using CHOPCHOP (https://chopchop.cbu.uib.no/). Briefly, 0.6 µg sgRNA (**Dataset S1**) was incubated with 3 µg SpCas9 or Cpf1 protein for 10 minutes at room temperature and electroporated into $2x10^5$ $H9^{NGN2;TMEM192\text{-}HA}$ cells using Neon transfection system (Thermo Fisher Scientific) and sorted into 96-well dishes containing 300 µL full growth medium (composition as described above). Single cells were allowed to grow into colonies and duplicated for multiplex sequencing. Genomic DNA samples were obtained by incubating cells in 30 µL PBND (50 mM KCl, 10 mM Tris-HCl, pH 8.3, 2.5 mM $MgCl_2$-$6H_2O$, 0.45% NP-40 and 0.45% Tween-20) with protease K (40 µg/ml) at 37°C for 5 min and heated to 55°C and 95°C for 30min and 15 min, respectively. The first round of PCR was performed to amplify the target region using gene-specific primers (**Dataset S1**) that contain partial Illumina adaptor sequences (i.e., Forward primer: 5'-ACACTCTTTCCCTACACGACGCTCTTCCGATCT$[n]_{18\text{-}22}$ -3', Reverse primer: 5'-GTGACTGGAGTTCAGACGTGTGCTCTTCCGATCT$[n]_{18\text{-}22}$ -3', $[n]_{18\text{-}22}$ represent gene specific sequences). The resulting PCR products with adapter-modified ends can be further amplified in the second round of PCR by universal primers containing attachment sites for the flow cell and index sequences (i.e., Forward primer: 5'-AATGATACGGCGACCACCGAGATCTACACTCTTTCCCTACACGACGCTCTTCCGATCT -3', Reverse primer: 5'- CAAGCAGAAGACGGCATACGAGAT-$[n]_8$-GTGACTGGAGTTCAG ACGTGTGCT -3', $[n]_8$ represents index sequences). The final PCR products were purified using QIAquick PCR purification kit (Qiagen, 28106). Sequencing was performed using Miseq Reagent kits v2 on Illumina Miseq following the denature and dilute libraries guide of Miseq system, and out-of-frame indels were identified using OutKnocker (www.OutKnocker.org) (8). Knockout candidates were confirmed by Western blot on whole cell lysates or by proteomics (**Dataset S1**) and individual deletion sequences are provided in **Dataset S1**. $HeLa^{TMEM192\text{-}HA}$ Control as well as Control, *NPC1*$^{-/-}$ and *NPC2*$^{-/-}$ $H9^{NGN2;TMEM192\text{-}HA}$ cells have been previously reported (3).

**Organelle IP from HeLa Cells**

For LysoIP from HeLa cells, we employed a previously reported method: dx.doi.org/10.17504/protocols.io.bw7hphj6. Briefly, HeLa cells (untagged, Control, and *ASAH1*$^{-/-}$) were seeded in three 10 cm dishes per replicate and harvested at 70–80 % confluency. For LysoIP, 65 µL of anti-HA magnetic bead slurry (Pierce) was used per replicate, and samples were incubated with beads at 4 °C with gentle rotation for 30 min. From each incubation, 10 µL of flow-through was collected for Western blot analysis. Anti-HA beads were washed twice with high-salt KPBSi (150 mM NaCl) and once with KPBSi. Samples were eluted with 120 µL 0.5 % NP-40 in KPBSi at 4°C with gentle rotation for 30 min. From each eluate, 20 µL was used for Western blot analysis, and 100 µL was further processed for TMTpro proteomics (see below).

**Lipidomics**

For lipidomics of whole-cell and isolated lysosomes from HeLa cells, 5x15cm dishes per genotype (untagged, Control and *ASAH1*$^{-/-}$) were used as input for LysoIP and 6 wells of a 6 well per genotype for whole-cell samples. Whole-cell samples were washed three times with 1xPBS, and harvested in ice-cold 1xPBS (+ protease inhibitor cocktail) by cell scraping. Cells were pelleted, the supernatant aspirated and snap frozen. LysoIP was performed as described above with the modification that after the IP, HA-beads were washed, snap frozen and shipped for lipidomic analysis.

For lipid extraction, whole cell or LysoIP samples were processed in ice-cold milli-Q water by snap-freezing (in liquid nitrogen)/thawing (using an ultrasound water bath for 3 min) repeatedly and finally extracted as described(9). Protein concentration in the lysates was quantified using the BCA assay, and approximately 50 µg of lysate was transferred to Pyrex glass tubes with PTFE-lined caps. For lipid extraction, 6 mL of ice-cold chloroform and methanol (2:1 v/v) and 1.5 mL of water were added to each sample. The tubes were thoroughly vortexed to ensure homogeneous mixing of polar and non-polar solvents. SPLAH internal standards (Avanti Product Number 330707-1EA) were added before extraction. The samples were centrifuged at 1000 rpm for 20 minutes at 4°C to separate the organic and aqueous phases. The lower organic phase was carefully pipetted into a new glass tube using a sterile glass pipette, avoiding the intermediate layer containing cell debris and precipitated proteins. The organic phase was then dried under a nitrogen stream until all solvents had evaporated. Finally, the samples were reconstituted in 100 µL of Isopropanol: Acetonitrile: Water (60:35:5) and stored at -80°C prior to analysis.

For ganglioside extraction, the aqueous layer was collected and dried under a gentle stream of nitrogen. The dried aqueous phase was reconstituted in 1 mL pure water and subsequently desalted by using Sola HRP SPE 30 mg/2 mL 96-well plate 1EA (Thermo Scientific #60509-001). Initially, the cartridges were cleaned 3 times with 1 mL of MeOH and equilibrated 3 times with water. Samples were loaded onto the column, washed 3 times with water and, finally, the gangliosides were eluted by three times 1 mL of MeOH. The eluate was dried under nitrogen flow and reconstituted in MeOH/H2O/CHCl3 (60:9:120, v/v/v).

For LC-MS/MS analysis, lipids were separated using ultra-high-performance liquid chromatography (UHPLC) coupled with tandem mass spectrometry (MS/MS). UHPLC analysis was conducted on a C30 reverse-phase column (Thermo Acclaim C30, 2.1 x 250 mm, 3 µm) maintained at 55°C and connected to a Vanquish Horizon UHPLC system, along with an OE240 Exactive Orbitrap MS (Thermo Fisher Scientific) equipped with a heated electrospray ionization probe. Each sample (5 µL) was analyzed in both positive and negative ionization modes. The mobile phase included 60:40 water: acetonitrile with 10 mM ammonium formate and 0.1% formic acid, while mobile phase B consisted of 90:10 isopropanol: acetonitrile with the same additives. The elution was performed with a gradient of 90 min; for 0–7 min, elution started with 40% B and increased to 55%; from 7 to 8 min, increased to 65% B; from 8 to 12 min, elution was maintained with 65% B; from 12 to 30 min, increased to 70% B; from 30 to 31 min, increased to 88% B; from 31 to 51 min, increased to 95% B; from 51 to 53 min, increased to 100% B; during 53 to 73 min, 100% B was maintained; from 73 to 73.1 min, solvent B was decreased to 40% and maintained for another 16.9 min for column re-equilibration. Flow rate of 0.2 mL/min, injection volume of 5 µL, and column temperature of 55°C. Mass spectrometer settings included an ion transfer tube temperature of 300°C, vaporizer temperature of 275°C, Orbitrap resolution of 120,000 for MS1 and 30,000 for MS2, RF lens at 70%, with a maximum injection time of 50 ms for MS1 and 54 ms for MS2. Positive and negative ion voltages were set at 3250 V and 2500 V, respectively. Gas flow rates included auxiliary gas at 10 units, sheath gas at 40 units, and sweep gas at 1 unit. High-energy collision dissociation (HCD) fragmentation was stepped at 15%, 25%, and 35%, and data-dependent tandem MS (ddMS2) ran with a cycle time of 1.5 s, an isolation window of 1 m/z, an intensity threshold of 1.0 x $10^4$, and a dynamic exclusion time of 2.5 s.

Full-scan mode with $ddMS^2$ was performed over an m/z range of 250-1700, with EASYIC™ (ThermoFischer) used for internal calibration. The raw data were processed and aligned with LipidSearch 5.1 (ThermoFischer, #OPTON 30879), using a precursor tolerance of 5 ppm and a product tolerance of 8 ppm. Further filtering and normalization were conducted using Lipidcruncher(10). Semi-targeted quantification was performed by normalizing the area under the curve (AUC) to the AUC of internal standards and further normalized with the total quantified protein level.

For LC-MS/MS analysis of gangliosides, samples were analyzed using a Vanquish UHPLC system (Thermo Scientific) coupled to an Orbitrap Exploris 240 mass spectrometer (Thermo Scientific, #BRE725535). Separation was achieved on a Kinetex HILIC column (Phenomenex, #00D-4461-AN; 2.6 µm, 100 x 2.1 mm). The mobile phase consisted of solvent A (acetonitrile with 0.2% v/v acetic acid) and solvent B (water containing 10 mM ammonium acetate, pH 6.1, adjusted with acetic acid). The column temperature was maintained at 50 °C. A gradient elution was employed at a constant flow rate of 0.6 mL / min: 12.3 % B at 0 min, a linear increase to 22.1% B from 1 to 15 min, followed by column equilibration at 12.3% B for 5 minutes. Mass spectrometry analysis was performed in heated electrospray ionization (HESI) mode under the following conditions: spray voltage at −4.5 kV, heated capillary temperature at 300°C, and vaporizer temperature at 250°C. The gas settings were as follows: sheath gas at 40 units, auxiliary gas at 5 units, and sweep gas at 1 unit. The ion transfer tube temperature was maintained at 300°C. For MS1 analysis, the Orbitrap resolution was set to 120,000 with a scan range of 700–1800 m/z, RF lens at 60%, and an AGC target set to standard. For MS2, the Orbitrap resolution was set to 30,000. Internal calibration was achieved using EASY-IC. After data search as above, quantification was achieved by normalizing the area under the curve to GM3-d5 standards, followed by further normalization to the amount of protein used in the preparation.

**Label-free nDIA proteomics of iNeurons and iDA neurons**

Protocols describing our DIA proteomics experiments can be found at: DOI: https://dx.doi.org/10.17504/protocols.io.eq2lyo43pgx9/v1. Control and LSD mutant cells were seeded on Geltrex-coated tissue culture plates and differentiated according to the methods stated above. For d30 timepoints, Control, *GRN*$^{-/-}$, *GBA1*$^{-/-}$, *ASAH1*$^{-/-}$, *SMPD1*$^{-/-}$ cells were seeded in triplicates in coated 12 well plates. For all other timepoints, cells were replated at day 16 in triplicates into coated 12 well plates. Cells were dissociated using 0.5 mM EDTA-PBS and washed in PBS, pelleted at 2000g for 5 min, the supernatant aspirated and pellet snap frozen and stored at -80°C.

Cell pellets were lysed in 140 µL 8M Urea and 100 mM Tris pH 8.0 by syringe with a 21G needle. Protein concentration was measured by protein BCA assay. 10 mM TCEP and 40 mM chloroacetamide were added for reduction and alkylation. LysC (FUJIFILM Wako) was added to samples in a 100:1 ratio (protein-to-LysC) and the samples were incubated on a rocker for 4 h at room temperature. The urea was diluted to 2 M with 100 mM Tris pH 8.0. Trypsin was added to samples in a 100:1 ratio (protein-to-trypsin) and the samples were incubated on a rocker overnight at room temperature. Digested peptides were desalted using Waters 25 mg Sep-Pak tC18 96-well Plates (Cat. No.: 186002319). The concentration of desalted peptides was measured by peptide BCA assay. Separation was performed on a C18 capillary column (40 cm length × 100 µm inner diameter) packed with Accucore resin (2.6 µm, 150 Å, Thermo Fisher Scientific) at 55 °C and an Vanquish Neo LC system (Thermo Scientific). Flow rate was set to 450 nL/min. Mobile phase A consisted of 0.125% formic acid in ACN/water (5:95, v/v). Mobile phase B consisted of 0.125% formic acid in ACN/water (95:5, v/v). When peptides were loaded on the column, mobile phase B increased from 5% to 35% over 32 minutes, increased to 99% over 2 minutes, and stayed at 99% for 6 minutes. Eluting peptides were analyzed by an Orbitrap Astral mass spectrometer (Thermo Scientific) using nDIA(11, 12) and analyzed by DIA-NN (v2.1.0)(13, 14)·(15). Methionine oxidation and N-terminal acetylation were enabled as variable modifications. Precursor m/z range was set to 380 to 980. Fragment ion m/z range was set to 110 to 2000. All other settings were kept default.

**TMTpro 18plex proteomics**

*Proteomic sample preparation.* Sample preparation of proteomic analysis of whole-cell extract from HeLa control and mutant lysates performed according to previously published studies(3, 5) (see protocol at dx.doi.org/10.17504/protocols.io.rm7vzjej4lx1/v1). Replicate cell cultures were grown and treated independently and are considered biological replicates in the context of TMT experiments. Cells were washed twice with 1xPBS and harvested on ice using a cell scraper in 1xPBS. Cells were pelleted via centrifugation for 5 minutes (5000*g*, 4°C), and washed with 1xPBS before resuspending in lysis buffer (Urea, 150 mM TRIS pH 7.4, 150mM NaCl, protease and phosphatase inhibitors added). After a 10 second sonication, and optional French-pressing through a G25 needle, lysed cells were pelleted and protein concentration of clarified sample determined using BCA kit (Thermo Fisher Scientific, 23227). 100 µg protein extract of each sample were incubated for 30 minutes @ 37°C with 5 mM TCEP for disulfide bond reduction with subsequent alkylation with 25 mM chloroacetamide for 10 minutes at RT with gentle shaking. Methanol-chloroform precipitation of samples was performed as follows: To each sample, 4 parts MeOH was added, vortexed, one part chloroform added, vortexed, and finally 3 parts water added. After vortexing, suspension was centrifugated for 2 minutes at 14000*g* and the aqueous phase around the protein precipitate removed using a loading tip. Peptides were washed twice with MeOH and resuspended in 200 mM EPPS, pH 8, and digested for 2 h with LysC (1:100) at 37°C, followed by Trypsin digestion (1:100) at 37°C overnight with gentle shaking.

*Tandem mass tag (TMT) labeling*. 50 µL of digested samples were labeled by adding 10 µL of TMT reagent (stock: 20 mg/ml in acetonitrile, ACN) together with 10 µL acetonitrile (final acetonitrile concentration of approximately 30% (v/v)) for 2 h at room temperature before quenching the reaction with hydroxylamine to a final concentration of 0.5% (v/v) for 15 minutes. The TMTpro-labeled samples were pooled together at a 1:1 ratio, resulting in consistent peptide amount across all channels. Pooled samples were vacuum centrifuged for 1 hour at room temperature to remove ACN, followed by reconstitution in 1% FA, samples were desalted using C18 solid-phase extraction (SPE) (200 mg, Sep-Pak, Waters) and vacuum centrifuged until near dryness. Our protocol for these steps can be found at dx.doi.org/10.17504/protocols.io.rm7vzjej4lx1/v1.

*Basic pH reverse phase HPLC*. Dried peptides were resuspended in 10 mM $NH_4HCO_3$ pH 8.0 and fractionated using basic pH reverse phase HPLC(16). Samples were offline fractionated into 96 fractions over a 90 minutes run by using an Agilent LC1260 with an Agilent 300 Extend C18 column (3.5 µm particles, 2.1 mm ID, and 250 mm in length) with mobile phase A containing 5% acetonitrile and 10 mM $NH_4HCO_3$in LC-MS grade $H_2O$, and mobile phase B containing 90% acetonitrile and 10 mM $NH_4HCO_3$ in LC-MS grade $H_2O$ (both pH 8.0). The 96 resulting fractions were then pooled in a non-continuous manner into 24 fractions(17). This set of 24 fraction was divided into 2x12 sets (even or odd numbers), acidified by addition of 1% Formic Acid (FA) and vacuum centrifuged until near dryness. One set (12 samples) was desalted via StageTip, dried and reconstituted in 10 µL 5% ACN, 5% FA before LC-MS/MS processing. Our protocol for these steps can be found at dx.doi.org/10.17504/protocols.io.rm7vzjej4lx1/v1.

*Mass spectrometry acquisition*. For TMTpro proteomics, data collection was performed on Orbitrap Ascend Tribid mass spectrometer (for HeLa LysoIP proteomics) (Thermo Fisher Scientific, San Jose, CA), coupled with a FAIMS Pro device and a Proxeon EASY-nLC1200 liquid chromatography (Thermo Scientific). 10% of resuspended samples were loaded on a 35 cm analytical column (100 mm inner diameter) packed with Accurcore150 resin (150 Å, 2.6 mm, Thermo Fisher Scientific, San Jose, CA) for LC-MS analysis. Peptide separation was performed with a gradient of acetonitrile (ACN, 0.1% FA) from 3-13% (0-83 minutes) and 13-28% (83-90 minutes) during a 90 min run. LC-MS/MS was combined with 3 optimized compensation voltages (CV) parameters on the FAIMS Pro Interface to reduce precursor ion interference(18). Data-dependent acquisition (DDA) was performed by selecting the most abundant precursors from each CV's (-40/-60/-80) MS1 scans for MS/MS over a 1.25 second duty cycle. The parameters for MS1 scans in the Orbitrap include a 400-1,600 m/z mass range at 60,000 resolution (at 200 Th) with $4 \times 10^5$ automatic gain control (AGC) (100%), and a maximum injection time (max IT) of 50 ms. Most abundant precursors (with 120 s dynamic exclusion +/- 10 ppm) were selected from MS1 scans, isolated using the quadrupole (0.6 Th isolation), fragmented with higher-energy collisional dissociation (HCD, 36% normalized collision energy), and subjected to MS/MS (MS2) in the Orbitrap detector at 50,000 resolution, 5x AGC, 110 – 200 m/z mass range, IT 86 ms and with 120 s dynamic exclusion +/- 10 ppm.

*Data processing*. Raw mass spectra were converted to mzXML, monoisotopic peaks reassigned using Monocle (19) and searched using Comet (20) against all canonical isoforms found in the Human reference proteome database (UniProt Swiss-Prot 2019-01; https://ftp.uniprot.org/pub/databases/uniprot/previous_major_releases/release-2019_01/)) as well as against sequences from commonly found contaminant proteins and reverse sequences of proteins as decoys, for target-decoy competition (21). For searches, a 50-ppm precursor ion tolerance and 0.02 Da product ion tolerance for ion trap MS/MS as well as trypsin endopeptidase specificity on C-terminal with 2 max. missed cleavages was set. Static modifications were set for carbamidomethylation of cysteine residues (+57.021 Da) and TMTpro labels on lysine residues and N-termini of peptides (+304.207 Da);

variable modification was set for oxidization of methionine residues (+15.995 Da). Peptide-spectrum matches (PSMs) were filtered at 2% false discovery rate (FDR) using linear discriminant analysis (Picked FDR method, based on XCorr, DeltaCn, missed cleavages, peptide length, precursor mass accuracy, fraction of matched product ions, charge state, and number of modifications per peptide (additionally restricting PSM Xcorr >1 and peptide length>6, and after a 2% protein FDR target filtering(22) PSM reporter ion intensities were quantified. Quantification was performed using a 0.003-Da window around the theoretical TMT-reporter m/z, and filtered on precursor isolation specificity of > 0.5 in the MS1 isolation window and the output was filtered using summed SNR across all TMT channels > 200. MSstatsTMT(23) was performed on peptides with >200 summed SNR across TMT channels. For each protein, the filtered peptide–spectrum match TMTpro raw intensities were summed and $log_2$ normalized to create protein quantification values (weighted average) and normalized to total TMT channel intensity across all quantified PSMs (adjusted to median total TMT intensity for the TMT channels) (24). $Log_2$ normalized summed protein reporter intensities were compared using a Student's t-test and p-values were corrected for multiple hypotheses using the Benjamini-Hochberg adjustment(25). Subcellular and functional annotations were based on previous published list of high confidence annotations (26), "high" & "very high" confidence, additional manual entries from (5, 27), AmiGO Pathway online tool and mitochondrial annotation was based on MitoCarta 3.0 (28). GO-enrichment was performed with ShinyGO (29) or topGO.

**Analysis of proteomic datasets**

Resulting files were analysed using R scripts, which is available on GitHub (https://github.com/sauerkrausi/neuroLSD) and readme files for analysis pipelines can be found at 10.5281/zenodo.17296003).

*HeLa LysoIP TMT Proteomics:* We analyzed LysoIP-TMT data from HeLa cells (untagged, $HeLa^{TMEM192\text{-}HA}$ control, *ASAH1*$^{-/-}$; in triplicates) quantified with MSstats. Results were transformed to −log10(p/q), filtered for finite log2FCs, and joined to a curated subcellular annotation matrix with hydrolase genes flagged. Row-z heatmaps were generated by genotype and fraction, and cluster-level patterns were visualized with violin plots. Organelle-specific co-enrichment was tested relative to lysosomal means, followed by Wilcoxon tests with Benjamini–Hochberg correction. Overlap structure was visualized by Venn diagrams, and enrichment of biological processes was assessed with topGO.

*LFQ-nDIA of iN and iDA neurons:* Whole-cell proteomes from iN and iDA neurons (day 30 and 50; in triplicates per neuron and genotype) were analyzed as follows:

*Proteomics Data Processing and Quality Control:* Replicate-level proteomics data from iN and iDA differentiated at day 30 and day 50 were imported and subjected to multi-level quality assessment. Median relative standard deviation (RSD) and protein identification counts were computed per neuron type and genotype to summarize data quality and coverage. Log2 fold-change matrices were generated for all genotype-neuron-day combinations, clustered hierarchically, and joined with subcellular annotations to identify compartment-level proteome patterns. Replicate correlation was computed using Pearson correlation across all pairwise combinations (rep1-rep2, rep1-rep3, rep2-rep3) stratified by genotype, neuron type, and differentiation day. Technical versus biological noise was assessed via F-ratio analysis (between-genotype variance / within-genotype variance) per subcellular annotation, computed across genes with complete triplication. Samples with F-ratios > 1 indicate biological signal exceeding technical noise. Principal component analysis (PCA) was performed on log2-scaled intensities across all samples and after QC filtering to visualize proteome structure by genotype, neuron type, and differentiation timepoint. Protein identification counts were quantified per replicate and summarized by genotype-neuron combination.

*Subcellular Annotation and Summarized Fold Change:* Gene-level quantified intensities were matched to a curated subcellular annotation matrix. For each gene, genotype, neuron type, and day, replicate-level $log_2$ fold-change values were calculated relative to control (ctrl) baseline. Quantified genes were stratified by annotation and visualized via annotation-level heatmaps and summarized statistics. Missing values were encoded as NA and handled downstream per analysis.

*Neuronal Identity Validation:* Neuronal differentiation quality and cell-type identity were confirmed through expression profiling of a curated set of neuronal developmental markers (For **Fig. S2*D***: BDNF, CAMK2B, CRTC1, DCX, JUN, MAP2, NCAM1, NEFH, NEFL, NEFM, NES, POU3F2, SLC17A7, SYN1, SYP, TUBB3, TH, BSN, SYNJ1, GAP43, SYN2, SYN3), plus pre/postsynaptic markers and dopaminergic lineage markers (tyrosine hydroxylase). Synaptic machinery components (v-ATPase, SNARE complexes,

SV docking machinery) were profiled across all genotypes via ranked log2 fold-change visualizations. Knockout efficiency for key lysosomal genes was verified via barplots and heatmap visualization.

*Developmental Time Course (Day 30 vs. Day 50):* Neuronal maturation was tracked via trajectories of developmental markers (pre/postsynaptic proteins, neuronal maturation markers) across day 30 and day 50 timepoints for control iN and iDA. $Log_2$ fold-change values were normalized to day 30 baseline and plotted to visualize maturation-dependent proteome remodeling. Annotation-level comparisons identified compartment-specific dynamics across differentiation stages.

*Annotation-Stratified Comparative Analysis:* Violin plots of $log_2$ fold-change values were generated for user-selected subcellular annotations, stratified by genotype and neuron type to visualize effect-size distributions. Complementary volcano plots highlighted proteins with largest fold changes and statistical significance (fold change threshold and q-value threshold = 0.05). Heatmaps of compartment-specific proteins were generated per genotype, including specialized visualizations for organelle-specific complexes (e.g., autophagy machinery, mitochondrial complexes).

*Organelle-Level Correlation Analysis:* Spearman rank correlations of log2 fold-change profiles were computed between pairs of user-selected subcellular annotations across all genotypes. Correlation matrices were visualized via heatmaps and ranked by genotype-level deviation from the group mean (impact scoring). Bubble plots and correlation scatter plots compared correlation patterns between iN and iDA, stratified by disease class and compartment. Impact scores were calculated as the L2 norm (Euclidean distance) of each genotype's correlation profile from the group median, thereby quantifying proteome-wide disruption magnitude of inter-organellar communication networks. Genotypes with high impact scores indicate severe rewiring of organellar cross-talk, while low scores suggest preserved compartmental relationships despite KO perturbation. Euler diagrams were generated to visualize compartment-level protein overlap across genotypes, identifying shared versus genotype-specific proteome signatures within each subcellular annotation.

*Disease-Class Focused Subset Analyses:* Sphingolipidosis samples were analyzed as a disease-class cohort. Annotation-level heatmaps and correlation matrices were computed for this subset, and compartment-specific vulnerability patterns were compared across iN/iDA and timepoints.

**Protein-Protein Interaction Analysis of Neuronal Proteome**

*PPI reference database.* Protein-protein interactions were based on two reference PPI datasets from the Baker and Walther labs (30, 31). A master database of human PPIs was assembled from the Human PPI dataset (80% expected precision, hereafter P80; ~29,000 edges) and the Human Predictome Top 16k set (~16,500 edges, ~90% precision). Pairwise interactions were matched by undirected UniProt ID pairs and merged from both sources. Approximately 7,000 edges were supported by both datasets (convergent evidence); ~22,000 were exclusive to P80 and ~9,500 to Predictomes. All prediction scores were retained from both sources (AFMprob, RFprob, CFprob from P80; SPOC, ipTM_max, KIRC from Predictomes), together with localization data and orthogonal database evidence (STRING, BioGRID, PDB exact structure or homology model) from P80. For additional complex-level confirmation, human entries from CORUM (32) and ComplexPortal (33) were integrated by gene name and UniProt ID respectively. Subcellular annotations were added to each protein node. **Dataset S4** contains all of the associated data described above.

*Orthogonal support score.* An additive support score (maximum 9) was computed per edge to reflect the breadth and strength of orthogonal evidence: PDB exact structure (+3), PDB homolog (+1), ComplexPortal strict dimer (+3), ComplexPortal relaxed (member of larger complex, +1), STRING (+1), BioGRID (+1), CORUM (+1).

*Confidence tiering.* Edges were stratified into three tiers. Tier 1 (high confidence): convergent evidence across both prediction sources, or AFMprob ≥ 0.9 and RFprob ≥ 0.9, or SPOC ≥ 0.95. Tier 2: AFMprob ≥ 0.8, RFprob ≥ 0.8, or SPOC ≥ 0.9. Tier 3: all remaining edges. Only Tier 1 PPIs were used for all downstream analyses.

*Neuronal PPI baseline.* The Tier 1 PPI set was filtered to interactions where both protein nodes were detected in the control proteome of either iNeuron or iDA neuron datasets, yielding the neuronal baseline network. Each protein node was additionally annotated with subcellular localization and organelle membership derived from a binary annotation matrix.

*Trimer and tetramer detection.* Trimers and tetramers were enumerated exhaustively on the neuronal baseline network using igraph. For each detected trimer and tetramer, per-edge evidence was retrieved from the baseline and an aggregate support score computed as the mean support score across all constituent edges (max 9 per edge). Each structure was then assigned an evidence validation tier in descending order of confidence: (i) all edges supported by exact PDB crystal structure, (ii) all edges present in CORUM or ComplexPortal, (iii) two of three edges with exact PDB, (iv) two of three edges in CORUM or ComplexPortal, (v) two of three edges with any PDB evidence, (vi) at least one edge with PDB, CORUM, or ComplexPortal support, (vii) no orthogonal database evidence (novel). All trimers and tetramers were independently cross-referenced against CORUM and ComplexPortal.

*Complex growing and merging.* Higher-order complexes were grown iteratively from trimer and tetramer seeds, similar to a previous report (31). At each growth step, candidate proteins connected by Tier 1 PPIs to at least two existing complex members were added; growth continued until no further candidates met this criterion or the complex reached a maximum size of 100 subunits. Complexes derived from different seeds were subsequently merged if their member overlap exceeded 80% (smaller absorbed into larger), yielding a non-redundant set of predicted complexes.

*Complex quality control.* Grown complexes were validated against CORUM and ComplexPortal using two criteria: strict match, defined as all predicted members co-occurring in a single reference complex; and relaxed match, defined as ≥50% of predicted members present in a single reference complex. Conversely, a top-down coverage analysis assessed what fraction of known CORUM and ComplexPortal complex subunits were detected in the neuronal baseline, tiered as complete (100%), high (≥80%), partial (≥50%), or low (<50% subunit coverage). This bidirectional comparison allowed classification of each detected structure as known or novel relative to curated databases.

*KO Vulnerability Analysis.* To assess PPI network vulnerability, proteins significantly reduced in each KO relative to control were identified separately for iNeuron and iDA neurons. Significance was determined by FDR-corrected q-values ($q < 0.05$, primary) with p-values retained for comparison. For each KO x neuron type combination, every Tier 1 baseline PPI was assigned a state: an edge was classified as "lost" if either interacting protein was significantly reduced, and "retained" if both proteins had quantitative data but neither met the loss threshold. Where an edge was lost, the loss driver was annotated as node A, node B, or both. Edges were additionally stratified by shared subcellular compartment to resolve organelle-level vulnerability. Trimer and tetramer vulnerability was assessed by flagging structures as "disrupted" if at least one member was significantly lost in a given KO x neuron context. For higher-order complexes, the disruption criterion was size-dependent: complexes of ≤4 members were flagged if at least one member was lost; for complexes of >4 members, the threshold was ≥25% of members lost. Complex vulnerability was visualized as a binary heatmap across all complexes and KOs, separately for iNeuron and iDA neurons, with complexes annotated by their origin (trimer- or tetramer-seeded) and KOs annotated by disease class.

**Light Microscopy**

*$Ca^{2+}$-imaging in iNeurons and iDA neurons using spinning disk microscopy.* iNeurons and iDA neurons were differentiated to the indicated time points in µ-Slide 8-well glass-bottom plates (ibidi, #80807). On the day of measurement, cells were incubated with Fluo-4 for 30 min at 37 °C, washed with warm PBS, and fresh ND2 medium was added. Neurons were imaged at 37 °C and 5% $CO_2$ using a Nikon Eclipse Ti2-E motorized spinning disk confocal microscope (as described above) with a Nikon Plan Lambda 20×/0.75 N.A. air objective in 2×2-pixel binning mode (785×570 px). For each time-lapse, 1000 frames were recorded at 10 frames/s (100 ms exposure at 5% 488 nm laser power). Calcium dynamics were recorded under basal conditions, stimulation with 200 mM KCl, or inhibition with 50 µM CNQX and 20 µM D-AP5. Images were analyzed using cellposeSAM and R with custom-trained models and scripts (see https://github.com/sauerkrausi/neuroLSD and 10.5281/zenodo.17296003). Time-lapses were segmented with custom ML models in cellposeSAM, and the resulting masks were matched to fluorescence intensity over time for each ROI. Traces were baseline-corrected ($\Delta F/F$), binarized, and evaluated for network synchrony and event dynamics. Batch processing enabled per-cell trace extraction and correlation analysis. A protocol can be found at: DOI: https://dx.doi.org/10.17504/protocols.io.j8nlkzrzwl5r/v1.

*Immunocytochemical analysis.* A protocol can be found at: DOI: https://dx.doi.org/10.17504/protocols.io.bp2l6jyj1vqe/v1. iNeurons at the indicated day of differentiation were fixed with warm 4 % paraformaldehyde (Electron Microscopy Science, #15710, purified, EM grade) in PBS at 37°C for 30 min and permeabilized with 0.5 % Triton X-100 in PBS for 15 min at room temperature. After three washes with 0.02% Tween20 in PBS (PBST), cells were blocked for 10 minutes in 3 % BSA-1xPBS at room temperature and washed again three times in PBST. Cells were incubated for 3 h in primary antibodies in 3 % BSA-1xPBS and washed three times with PBST. Antibodies used include α-MAP2, α-

TH, α-SYN1, α-tubulin, α-NEFH, α-BSN, and α-SYP Secondary antibodies (Thermo Scientific, 1:400 in 3 % BSA-1xPBS) where applied for 1 hour at room temperature. To stain nuclei, Hoechst33342 (1:10000) was added for 5 minutes to cells in PBST and finally washed three times.

*Fixed-cell microscopy – general acquisition parameters.* A protocol can be found at: DOI: https://dx.doi.org/10.17504/protocols.io.bp2l6jyj1vqe/v1. Immunofluorescently labelled cells were imaged at room temperature using a Yokogawa CSU-W1 spinning disk confocal on a Nikon Eclipse Ti2-E motorized microscope equipped with a Nikon Plan Apochromat 40×/0.40 N.A air-objective lens, Nikon Plan Apochromat 60×/1.42 N.A oil-objective lens and a Plan Apochromat 100×/1.45 N.A oil-objective lens. Signals of 405/488/568/647 fluorophores were excited in sequential manner with a Nikon LUN-F XL solid state laser combiner ([laser line – laser power]: 405 nm - 80 mW, 488 nm - 80 mW, 561nm - 65 mW, 640 nm - 60 mW using a Semrock Di01-T405/488/568/647 dichroic mirror. Fluorescence emissions were collected with Chroma ET455/50m [405 nm], 488 Chroma ET525/50m [488 nm], 568 Chroma ET605/52m [561 nm], 633 Chroma ET705/72m [640 nm] filters, respectively (Chroma Technologies). Confocal images were acquired with a Hamamatsu ORCA-Fusion BT CMOS camera (6.5 $\mu m^2$ photodiode, 16-bit) camera and NIS-Elements image acquisition software. Consistent laser intensity and exposure time were applied to all the samples, and brightness and contrast were adjusted equally by applying the same minimum and maximum display values in ImageJ/FiJi (34).

*Evaluation of TH positive neurons.* A protocol can be found at: DOI: https://dx.doi.org/10.17504/protocols.io.bp2l6jyj1vqe/v1. Time-lapse calcium imaging data in ND2 format were processed using a custom Python pipeline implementing CellposeSAM segmentation. Image stacks were read with the nd2 package, and maximum intensity projections were generated prior to segmentation with a pretrained Cellpose model optimized for neuronal soma. Segmentation masks were filtered by area (100–10,000 pixels) to exclude artifacts and outliers. For each valid region of interest, centroid coordinates were extracted, and mean fluorescence intensities were computed across all time frames. Segmentation results were saved as 16-bit TIFF masks and visualization overlays, while per-cell centroid coordinates and intensity traces were exported as CSV files for downstream analysis and plotting in R.

Evaluation of synaptic proteins in iN and iDA samples. A protocol can be found at: DOI: https://dx.doi.org/10.17504/protocols.io.bp2l6jyj1vqe/v1. The quantitative measurement of select synaptic markers (for example, SYP, Bassoon) in iN and iDA neurons was performed using a custom Python pipeline. Maximum-intensity projection images were parsed for experimental metadata, and cytoskeletal channels were used to generate neuronal masks. Marker-specific background subtraction and segmentation routines were applied to detect synaptic puncta, followed by measurement of object morphology and fluorescence intensity. Colocalization between SYP and Bassoon was evaluated at the object level, and results were exported as cleaned images, QC overlays, and CSV files containing per-object and per-image metrics for downstream analysis.

**Electron Microscopy**

iNeuron and iDA cells of Control and select mutant cells were grown on Aclar plastic coverslips in above stated growth conditions until 70-80 % confluency was reached, washed twice in 1x PBS and fixed with a fixation mixture of 2 % formaldehyde and 2.5 % glutaraldehyde in 0.1 M Sodium Cacodylate buffer, pH 7.4 for 1 hour at room temperature. Sample preparation and microscopy was performed by the Harvard Medical School Electron microscopy facility (https://electron-microscopy.hms.harvard.edu/).

**Cryo-ET**

Associated protocols can be found on protocols.io (https://www.protocols.io/view/sample-preparation-and-vitrification-of-cell-cultu-yxmvme659g3p/v1, https://www.protocols.io/view/cryo-plasma-focused-ion-beam-pfib-milling-e6nvw1rozlmk/v1, https://www.protocols.io/view/cryo-et-data-acquisition-tomogram-reconstruction-a-dm6gpznxdlzp/v1).

*Sample preparation and freezing.* HeLa$^{TMEM192\text{-}HA}$ Control and *ASAH1*$^{-/-}$ cells (3) were cultured on EM grids as follows: 200-mesh gold grids with Silicon Dioxide R1/4 film (Quantifoil) were plasma cleaned, coated by incubation with 0.25 µg/mL laminin (Sigma-Aldrich, L4544) for 1h under the UV light of a sterile environment and afterwards washed twice with PBS. One day before plunging, laminin-coated grids are transferred to 8-well µ-Slide dishes (Ibidi, 80826) and ~12,000 cells are seeded per well together with 0.1mg/mL 10kDa Dextran coupled to Alexa Fluor 647 (Thermo Fisher Scientific, D22914) for endolysosome detection under cryogenic conditions. The next day, grids were vitrified with 3.5 µL of PBS-diluted autofluorescent 1 µm diameter Dynabeads (Thermo Fisher Scientific, acid65011) after 6.5s blotting time at

37°C and 70% relative humidity in liquid ethane (at -184°C) using an EM GP2 plunger (Leica Microsystems). After plunging, the grids were clipped into autogrids with a cutout for FIB-milling.

Cryo-Focused ion beam (FIB) milling. TEM-transparent lamellae were produced in a commercially available Aquilos2 dual-beam cryo-Focused Ion Beam and scanning electron microscope (cryo-FIB/SEM) instrument (Thermo Fisher Scientific). Intracellular lysosome fluorescence was observed using the integrated fluorescence light microscope (iFLM) of the Aquilos system. Fluorescent stacks were acquired using the 470 nm and 625 nm channels (4% laser intensity, 20ms exposure per slice, z-stack height of 6 µm, 15 slices). Correlation of the fiducial beads from the fluorescence 2D projection image with the grid's respective SEM overview image was carried out in MAPS v.3.28 (Thermo Fisher Scientific).

Semi-automatic FIB-milling was done using AutoTEM v.2.4.2 (Thermo Fisher Scientific) at a milling angle of 8°, as described (35). In short, the milling consisted of the following steps: i) rough milling at 1 nA to 1 µm thickness, (ii) 0.5 nA to 750 nm, and (iii) 0.3 nA to the 500 nm. Afterwards, the lamella was polished at a current of 50 pA to 150 nm) and 10 pA to 130 nm (with 0.2° overtilting). In some cases, remnants of the cell top surface with its organometallic layer had to be removed in addition to make the full tilt range in cryo-ET accessible.

Cryo-ET data acquisition and Processing. TEM data acquisition was performed on a Krios G4 at 300 kV with Selectris X energy filter and Falcon 4i camera (Thermo Fisher Scientific) using SerialEM v.4.1 and v.4.2(36). Tilt series were acquired at a nominal magnification of 64,000X (pixel size 1.971 Å) with a 10 eV energy slit using a dose-symmetric tilt scheme with an angular increment of 2°, a dose of ~2.4 $e^-$/$Å^2$ per tilt resulting in ~150 e/$A^2$ for the 61 tilt of the complete tilt series, and a target defocus between -3 and -5 µm. Tilt series were collected ranging from -52° to +68° relative to the lamella pretilt. The positions for tilt series acquisition were determined by visual inspection of 8,700X magnification lamella montage maps. 10 frames per tilt were acquired and aligned in SerialEM during acquisition.

Tomogram reconstruction and segmentation. The tilt series mrc stacks were corrected for dose exposure and projections of low-quality were removed manually. The tilt series was aligned using patch-tracking and reconstructed at bin4 using AreTomo (v.1.0.0) (37) and IMOD (v.4.12.62)(38). Tomogram denoising at bin4 (7.884Å/px) was done using IsoNet2 (39) or CryoCare as indicated (40). All membranes in the tomograms were segmented with Membrain-Seg (https://github.com/teamtomo/membrain-seg) (41) using the publicly available pretrained model (v10). For visualization purposes, all segmentations were manually touched up in Napari (10.5281/zenodo.17296003) and rendered using ChimeraX v.1.10 (42).

**Software and Resources**

The following software, packages, and resources were used for data analysis, visualization, and figure preparation:

R Environment: R (v4.5.2; R Project for Statistical Computing, RRID:SCR_001905); RStudio (v2026.01.1+403; Posit Software, RRID:SCR_000432); bigstatsr (1.6.1); broom (1.0.12, RRID:SCR_026874); Cairo (1.6.2); CalNetExploreR (0.1.0); car (3.1-2, RRID:SCR_022137); circlize (0.4.16, RRID:SCR_002141); ComplexHeatmap (2.25.2, RRID:SCR_017270); ComplexUpset (1.3.3, RRID:SCR_022752); corrplot (0.95, RRID:SCR_023081); cowplot (1.2.0, RRID:SCR_018081); data.table (1.18.2.1, RRID:SCR_026117); devtools (2.4.5, RRID:SCR_016961); dplyr (1.2.0, RRID:SCR_016708); factoextra (1.0.7, RRID:SCR_016692); fmsb (0.7.6); forcats (1.0.1); furrr (0.3.1); ggbiplot (0.6.2, RRID:SCR_025581); ggdendro (0.2.0); ggdist (3.3.3); gghalves (0.1.4); gghighlight (0.5.0); ggplot2 (4.0.2, RRID:SCR_014601); ggpp (0.6.0); ggpmisc (0.6.3); ggpubr (0.6.3); ggraph (2.2.1, RRID:SCR_021239); ggrepel (0.9.7, RRID:SCR_017393); ggsci (2.9); ggsignif (0.6.4, RRID:SCR_023047); ggVennDiagram (1.5.7, RRID:SCR_026950); grid (4.5.0); gridExtra (2.3, RRID:SCR_025249); igraph (2.2.2, RRID:SCR_019225); irlba (2.3.5.1); limma (3.64.1, RRID:SCR_010943); lintr (3.2.0); lubridate (1.9.5); magick (2.8.7); msstats (4.10.0, RRID:SCR_014353); NatParksPalettes (0.2.0); org.Hs.eg.db (3.21.0, RRID:SCR_024739); patchwork (1.3.2, RRID:SCR_024826); pheatmap (1.0.13, RRID:SCR_016418); plotly (4.11.0, RRID:SCR_013991); plyr (1.8.9, RRID:SCR_026985); png (0.1.8); qs (0.27.3); purrr (1.2.1, RRID:SCR_021267); RColorBrewer (1.1.3, RRID:SCR_016697); readr (2.2.0); reshape2 (1.4.4, RRID:SCR_022679); rlang (1.1.7); scales (1.4.0, RRID:SCR_019295); signal (1.8.1); stringr (1.6.0, RRID:SCR_022813); superheat (0.1.0); Ternary (2.3.6); tidyplots (0.3.1); tidyverse (2.0.0, RRID:SCR_019186); tidyr (1.3.2, RRID:SCR_017102); tibble (3.3.1, RRID:SCR_026493); timecourse (1.68.0, RRID:SCR_000077); topGO (2.60.1, RRID:SCR_014798); umap (0.2.10.0); UpSetR (1.4.0, RRID:SCR_026112); uwot (0.2.3); viridis (0.6.5, RRID:SCR_016696); zoo (1.8.14). Gene Ontology enrichment was performed using ShinyGO (Ge et al., 2020, RRID:SCR_019213).

Python Environment: Python (v3.12.2, RRID:SCR_008394); cellpose (≥4.0.5, RRID:SCR_021716); cellposeSAM (v4.0.6); matplotlib (latest version, RRID:SCR_008624); nd2 (latest version); numpy (latest version, RRID:SCR_008633); pandas (latest version, RRID:SCR_018214); pyqt6, qtpy, pyqt6-sip, pyqtgraph, superqt (latest versions); scikit-image (latest version, RRID:SCR_021142); tifffile (latest version, RRID:SCR_023338); tqdm (latest version, RRID:SCR_025100); torch (≥2.7.0, RRID:SCR_018536); torchvision (≥0.22.0).
Other software/databases: AmiGO (RRID:SCR_002143), CellProfiler (v4.2.1, RRID:SCR_007358), Comet 2019.01 (RRID:SCR_011925), CUDA (RRID:SCR_013225), DIA-NN (RRID:SCR_022865), FiJi/ImageJ 1.53t30 (RRID:SCR_002285), Fragpipe (RRID:SCR_022864), Lipidcruncher (10), LipidSearch 5.1 (RRID:SCR_023716), MassPike (43), Monocle (RRID:SCR_018685), NIS Elements 5.21.3 (RRID:SCR_014329), SEQUEST (20).

Artificial Intelligence Software: Code development, documentation, and debugging, was assisted by Claude and Cowork (Anthropic; models claude-sonnet-4-6 and claude-opus-4-6).

### Supplemental Figures

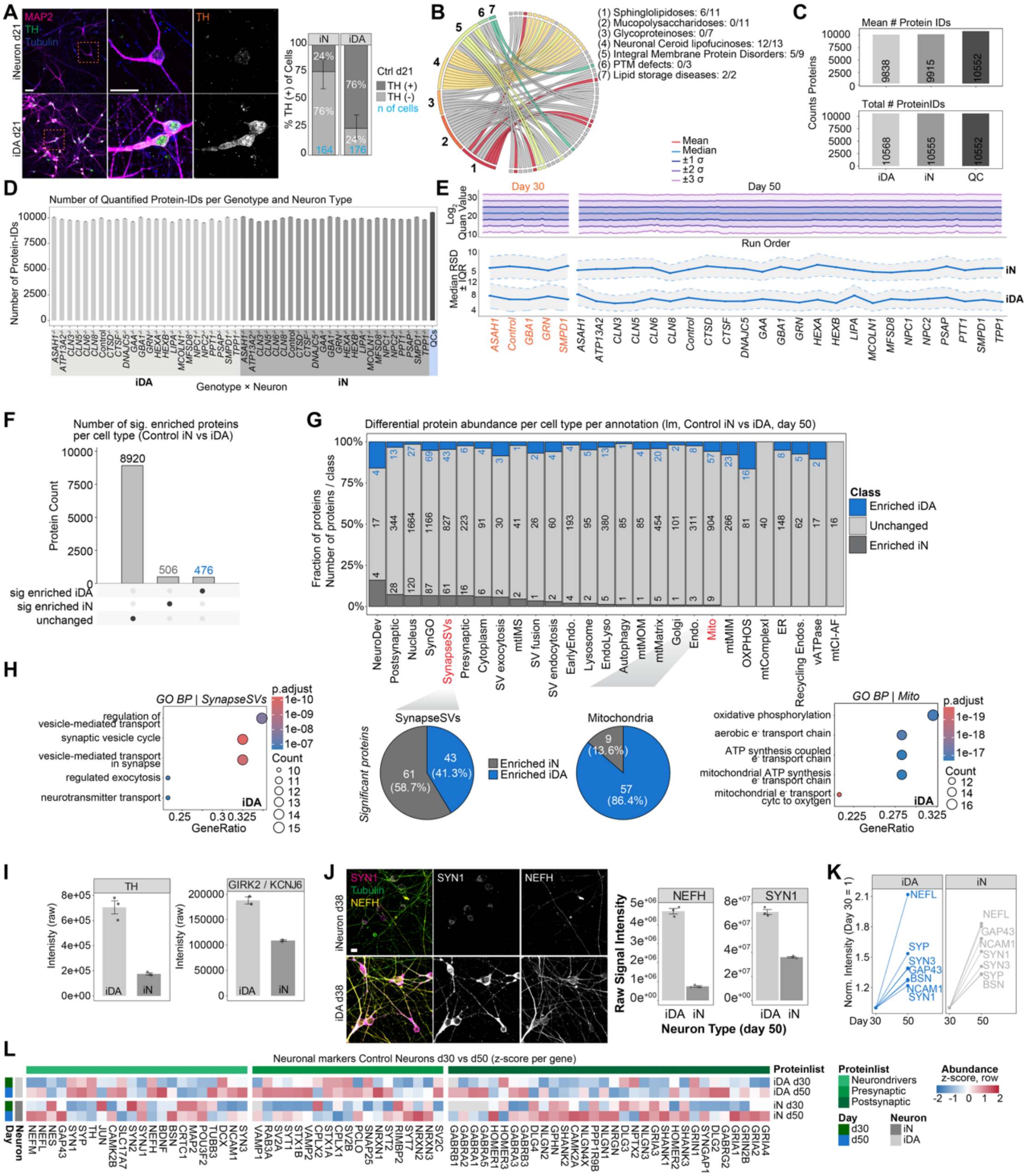


**Fig. S1. Proteome atlas of LSD neuronal cell models.** (***A***) Left: Confocal images of Control iN and iDA neurons at day 21 of *in vitro* differentiation, labeled for MAP2 (magenta), TH (green) and Tubulin (blue). Scale bar: 10 µm. Right: evaluation of % TH (+) neurons in iN or iDA (n(cells): iN = 164; iDA = 176). (***B***) Circos plot of LSD deletion coverage in human ES cells. LSD gene editing campaign focused on three disease-classes (Sphingolipidoses, NCL and Integral Membrane Protein Disorders). (***C***) Mean and total number of quantified IDs per sample across neuron types for day 50. (***D***) Histogram of total number of quantified proteins across genotypes and cell types. (***E***) Top: Lineplot of mean, median and sigma (range: 1-3) of $\log_2$ quant value across run order for day 30 and day 50 cohort. Bottom: As above, but plotted for median RSD ± IQR for iN (top) and iDA (bottom). (***F***) Analysis of the number of proteins enriched in either iDA or iN cells where $\log_2$FC>1.0 and p-value<0.05. (***G***) Distribution of differentially expressed proteins in iDA (blue) or iN (dark gray) cells across organelle or protein complex annotations. (***H***) Gene Ontology (GO)

analysis of synapseSV proteins or mitochondrial proteins enriched in iDA cells. The pie charts (center) indicate the number of proteins and percent of total for each compartment. (***I***) Raw $MS^1$ intensities for TH and GIRK2 proteins in Control iN and iDA cells. (***J***) Left: Confocal images of Control iN and iDA neurons at day 38 of differentiation, immunostained for SYN1 (magenta), NEFH (yellow) and Tubulin (green). Scale bar: 10 μm. Right: Raw $MS^1$ signal intensities for proteomic evaluation of SYN1 and NEFH in iN and iDA cells at day 50 of differentiation. (***K***) Line plot depicting intensity of select neurogenesis markers on day 30 and day 50 in iN and iDA neurons from Control cells (normalized to day 30). (***L***) Relative abundance of neuronal drivers, pre-synaptic, and post-synaptic proteins iN and iDA cells as measured by $MS^1$ intensities. Error bars, standard deviation.

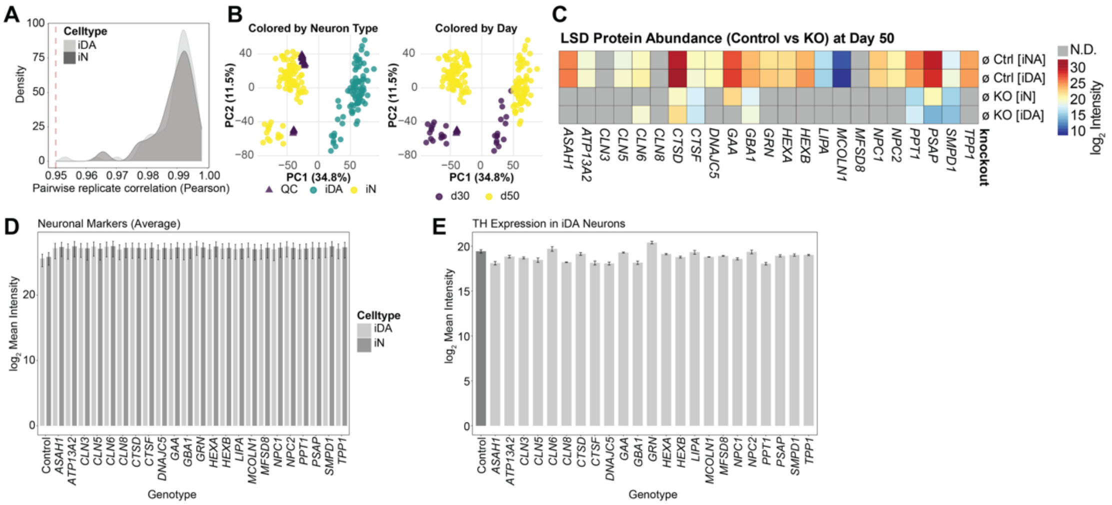


**Fig. S2. Data quality analysis and differentiation across LSD neuronal cell models.** (***A***) Plot of pairwise replicate Pearson correlation for iN and iDA cells. (***B***) PCA plots of total proteome datasets, color by neuron type and day of differentiation for iDA and iN cells. QC, refers to quality control standards included during the extended data collection period on the mass spectrometer. (***C***) Heatmap of target LSD protein abundance in Control and deletion mutants for iN and iDA. **See SI Appendix, Table S1**. (***D***) Log$_2$ mean MS$^1$ intensities for a set of neuronal marker proteins across Control and LSD mutant iN and iDA cells. See **METHODS** for list of proteins in category. Error bars, standard deviation. (***E***) Log$_2$ mean MS$^1$ intensities for the TH protein across Control and LSD mutant iDA cells. Error bars, standard deviation.

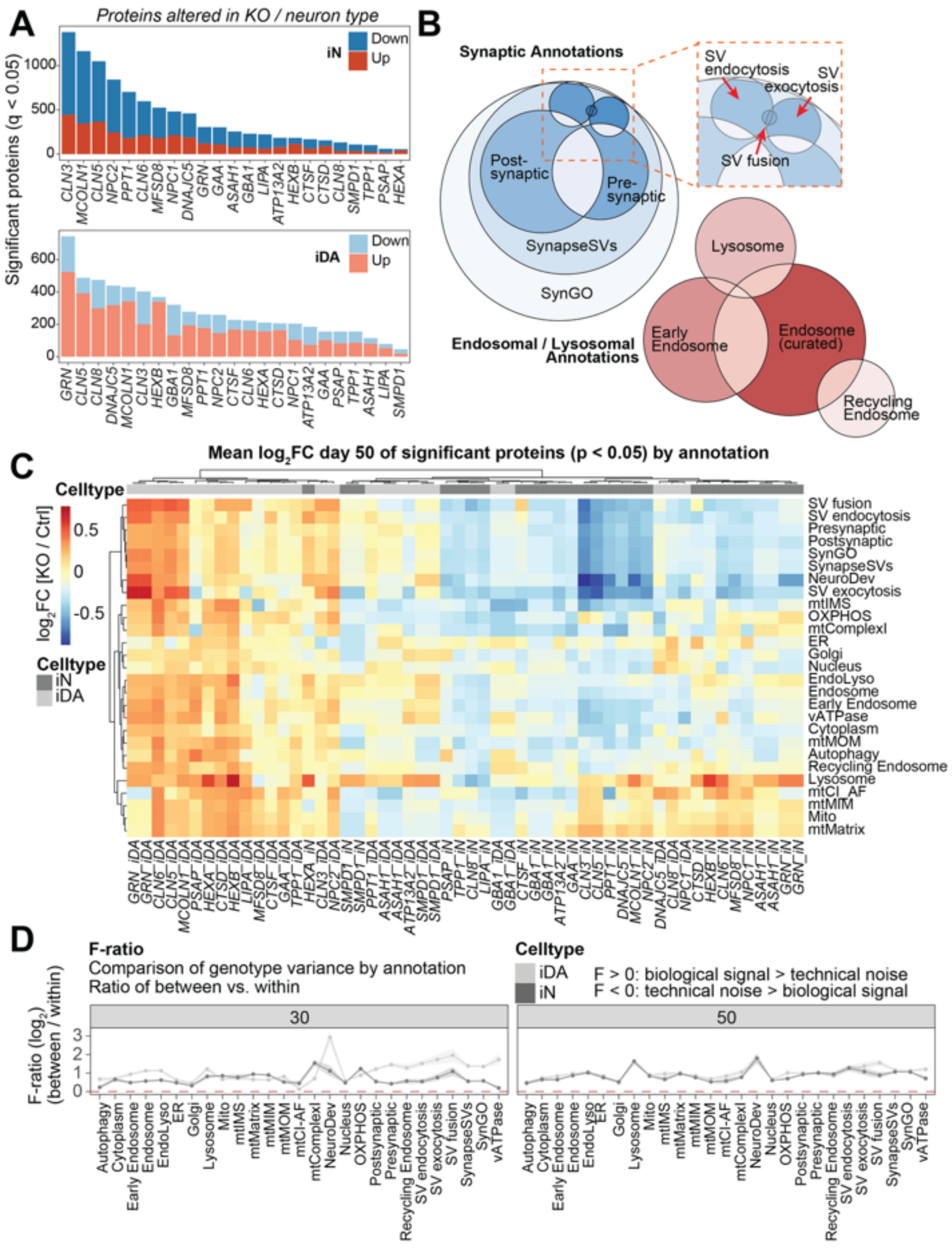


**Fig. S3. Expansion of endolysosomal and synaptic systems annotation across LSD iN and iDA neurons.** (***A***) Bargraph (descending order) depicting the number of proteins whose abundance is significantly (q-value < 0.05, FDR-corrected) increased (red shade) or decreased (blue shade) in the indicated LSD mutant iN or iDA cells compared to control neurons in same cell type. (***B***) Eulerplot depicting the overlap and scope of synaptic and endosomal/lysosomal annotations used for this study. Proteins in each category are listed in **SI Appendix, Dataset S1**. (***C***) Mean log$_2$FC of significantly altered proteins for individual organelles or protein complexes across LSD mutant iN or iDA cells. p-value<0.05. (***D***) Plot of F-ratio for variance by annotation ratio between and within genotypes. F-ratios > 0 have biological signals > technical noise. F-ratios < 0 have technical noise > biological signals.

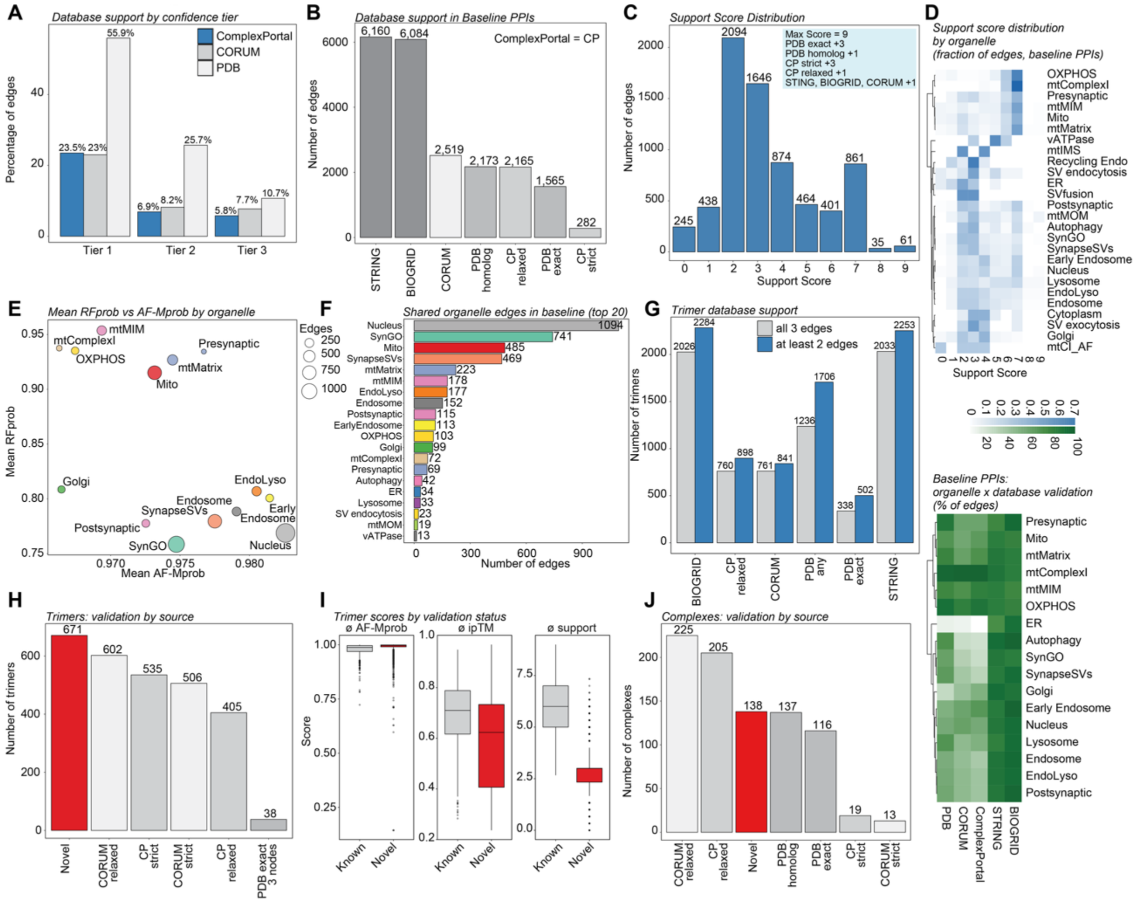


**Fig. S4. Establishment and QC of baseline protein interaction dataset for structural investigation of LSD mutant vulnerability analysis.** (***A***) Bargraph depicting structural database support (ComplexPortal, CORUM, PDB) for all PPI edges in tier 1-3 of PPIs. (***B***) Barplot of number of edges per bin of orthogonal validation information in neuronal baseline PPI dataset. (***C***) Histogram of cumulative support score distribution per PPI in neuronal baseline PPI dataset. (***D***) Heatmaps of average support score per PPIs per annotation (top) and fraction of PPIs found in validation databases expressed compared to all edges in annotation (bottom) in neuronal baseline PPI dataset. (***E***) Scatterplot of mean RFprob vs AF-Mprob per annotation / organelle in neuronal baseline PPI dataset. (***F***) Sorted bargraph depicting number of organelle edges per annotation / organelle in baseline PPI dataset (in descending order). (***G***) Barplot of number of trimer edges (grey = all three edges; blue = at least 2 edges) found in validation databases in baseline PPI dataset. (***H***) Bargraph of number of trimers found in validation databases in neuronal baseline PPI dataset. (***I***) Boxplots average scores of known and novel trimers for average AF-M probability (left), average ipTM score (middle) and cumulative support score (right) in neuronal baseline PPI dataset. (***J***) Bargraph of number of complexes found in validation databases in neuronal baseline PPI dataset.

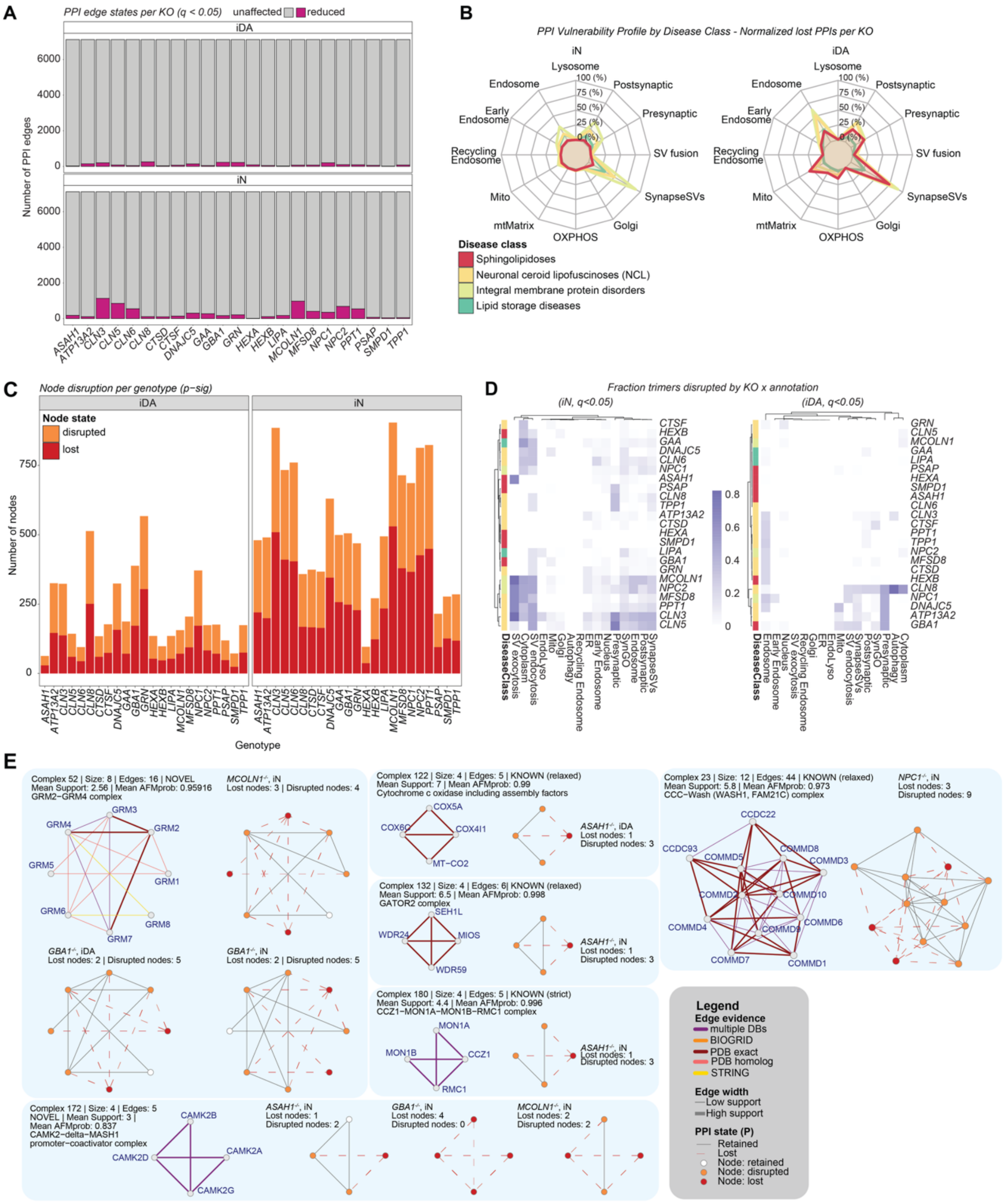


**Fig. S5. Analysis of pairwise PPIs and protein complex vlunerabilities across LSD mutant proteomes in iN and iDA cells.** (***A***) Stacked bargraph showing the number of unaffected and reduced (q < 0.05, FDR corrected) pairwise PPIs across LSD mutants for iDA (top) and iN (bottom) cells. (***B***) Radarplot showing PPI vulnerability by disease class for iN (left) and iDA (right) for 12 selected annotations / organelles. (***C***) Summary of node alterations [disrupted (orange) or lost (red)] across all LSD mutants for iN (right) and iDA (left) cells. (***D***) Heatmap depicting fraction of disrupted trimers per LSD mutant per annotation / organelle for iN (left) and iDA (right) cells. (***E***) Examples of complexes with lost or disrupted nodes include the glutamate receptor complex, CAMK2, cytochrome C oxidase, the GATOR complex, the CCZ1-MON1-RMC1 complex, and the Commander complex. The legend summarizes color codes for edge evidence and PPI state.

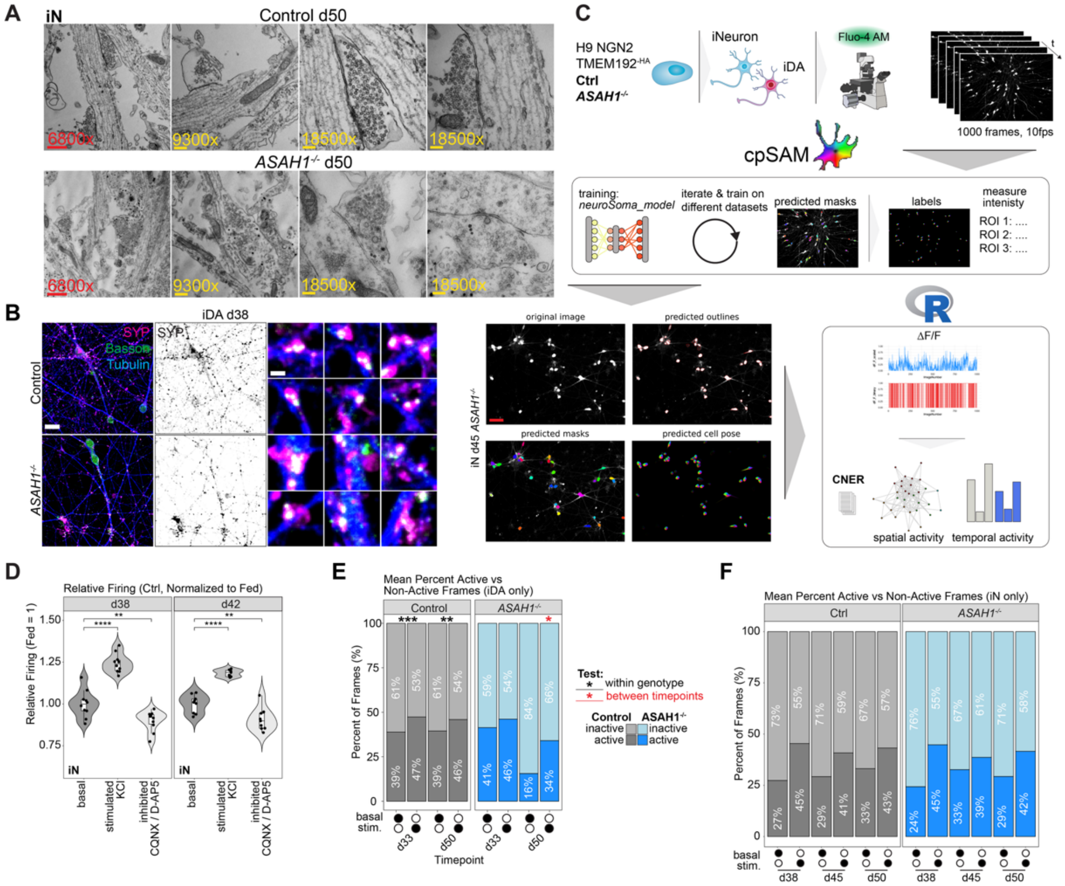

**Fig. S6. Synaptic morphology defects correlate with reduced basal Ca2+ firing activity in *ASAH1*$^{-/-}$ iDA cells.** (***A***) Representative EM images depicting synapses and neuronal ultrastructure of Control and *ASAH1*$^{-/-}$ iN cells at day 50 of *in vitro* differentiation. (***B***) Confocal images of Control and *ASAH1*$^{-/-}$ iN and iDA cells at day 38 of *in vitro* differentiation, labeled for SYP, Bassoon and Tubulin. Scale bar: 20 µm and 2 µm. The zoom-ins on the right are duplicated from **Fig. 5*B*.** (**C**) Schematic of Machine Learning-assisted neuronal calcium evaluation pipeline. nd2 time-series are segmented using custom-trained cellposeSAM (cpSAM) models. Calcium intensities over time were then analyzed in R. Stills of live-cell spinning-disk confocal images of day 45 iN cells (original image), predicted outlines and masks as well labels from cellposeSAM. Scale bar: 50 µm. (***D***) Violin plots of relative firing rate of day 38 and day 42 Control iN cells under basal, stimulated and inhibited conditions. d38: **P(0.006); ****P(0.0001); d42: **P(0.01);****P(0.0002). (***E***) Stacked bar graph for Control (left) and *ASAH1*$^{-/-}$ (right) iDA cells depicting the mean % of active and non-active frames (ROIs) for any given time. Data from day 33 and 50 of *in vitro* differentiation with or without stimulation by KCl. Control d33 fed vs KCL***P(0.0001); d50 fed vs KCL:**P(0.007); *ASAH1*$^{-/-}$ d33: fed vs KCL***P(0.00006); d33 vs 50 KCL: *P(0.03). (***F***) Stacked bar graph for Control (left) and *ASAH1*$^{-/-}$ (right) iN cells depicting the raw mean % of active ROIs at any given frame. Data from day 38, 42 and 50 of *in vitro* differentiation.

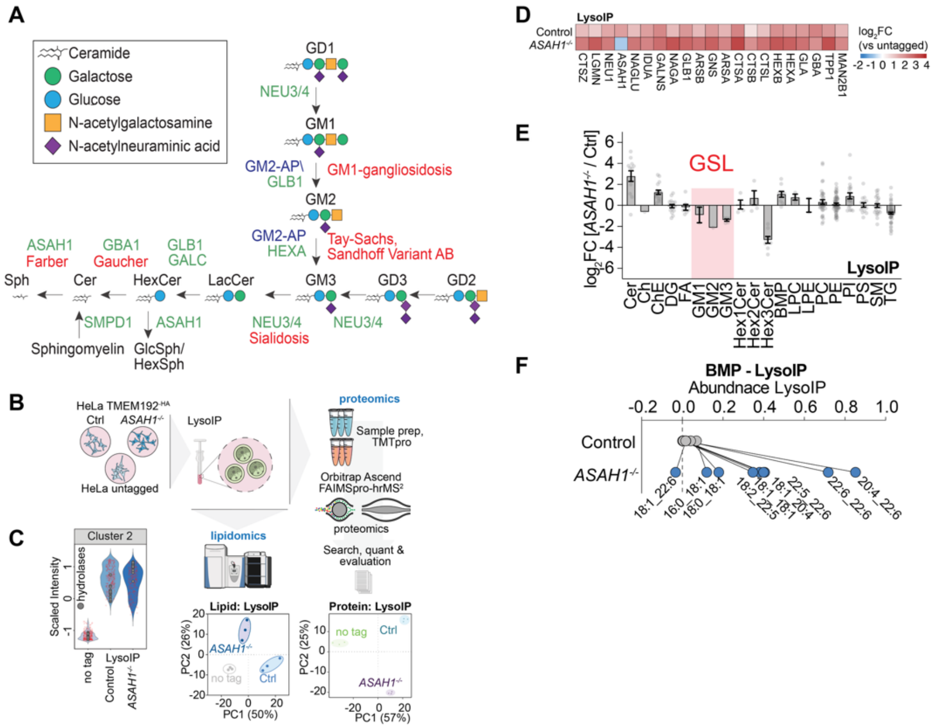


**Fig. S7:**

**Loss of *ASAH1*$^{-/-}$ disturbs lysosomal lipidome and endo-lysosomal morphology.** (***A***) Scheme depicting the pathway for Sphingolipid catabolism, including steps involving ASAH1, which converts ceramide to sphingosine and fatty acids. Modified from our previous publication (44). (***B***) Schematic for TMTpro-proteomics and label-free lipidomic of LysoIP samples from *ASAH1*$^{-/-}$, Control and untagged HeLa cells. PCA plots are shown below. (***C***) Violin plot of cluster 2 of heatmap (see **SI Appendix, Dataset S5**) depicting enrichment of proteins of Control and *ASAH1*$^{-/-}$ over untagged cells. Lysosomal hydrolases are shown as grey circles. (***D***) $Log_2FC$ of isolated organelle proteome and selected lysosomal hydrolases based on LysoIP proteomics of Control or *ASAH1*$^{-/-}$ cells vs untagged cells. TMTpro proteomics, based on n=3 biological replicates per genotype. (***E***) $Log_2FC$ of *ASAH1*$^{-/-}$ vs Control LysoIP lipidomics from HeLa cells. Data based on n=3 biological replicates per genotype. (***F***) Lineplot of select BMP species in Control and *ASAH1*$^{-/-}$ LysoIP samples. Data based on n=3 biological replicates per genotype.

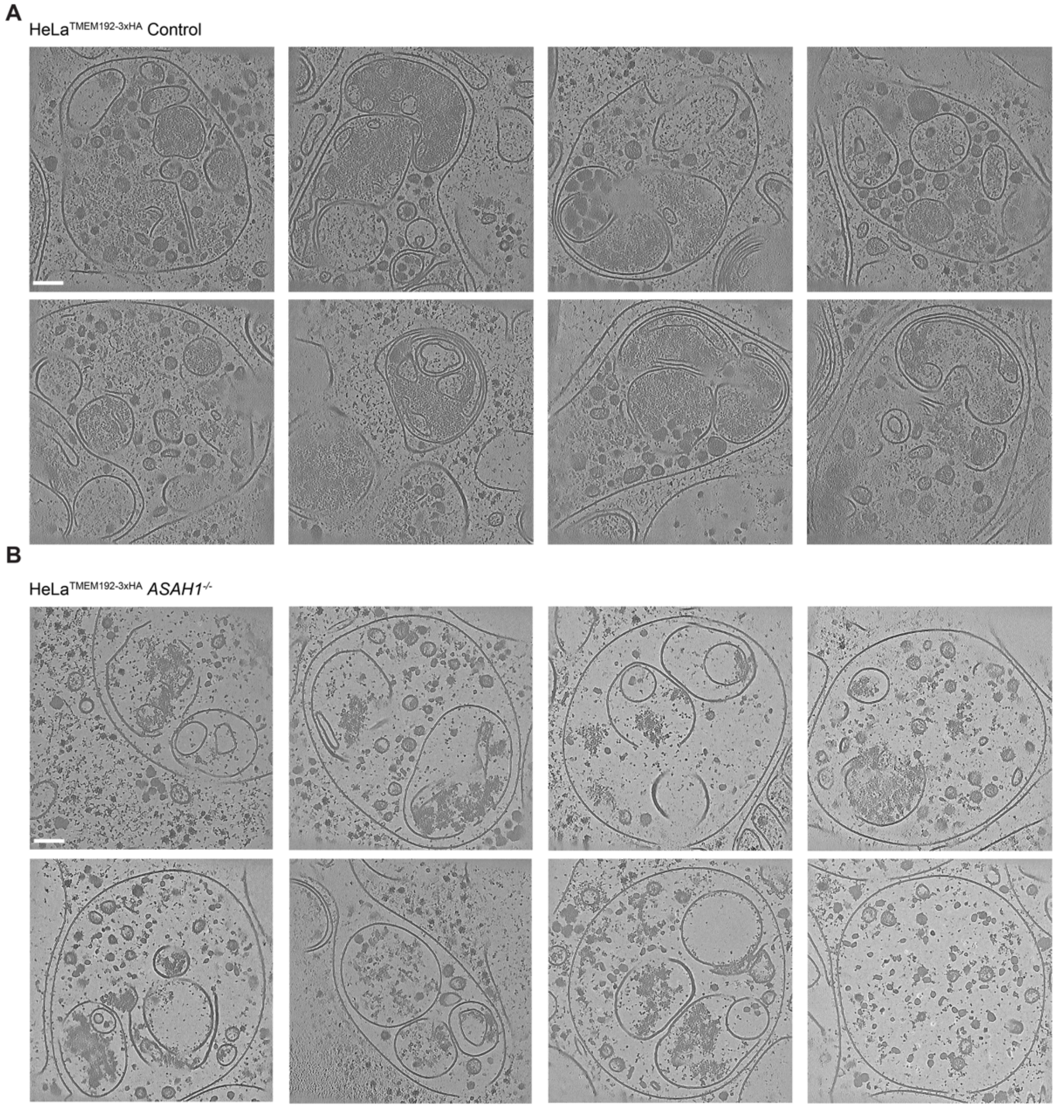


**Fig. S8: Analysis of Control and *ASAH1*$^{-/-}$ HeLa cells by cryo-ET. (*A*)** Gallery of representative tomograms of HeLa Control lysosomes. Scale bar 100 nm. **(*B*)** Gallery of representative tomograms of Hela *ASAH1*$^{-/-}$ lysosomes. Scale bar 100 nm. Images were denoised with IsoNet2 (39).

**Supplemental data legends**

**Movie S1 (separate file)**: Tomogram of endolysosome in Control HeLa cells, corresponding to **Fig. 5F (Top)**.

**Movie S2 (separate file)**: Tomogram of endolysosome in *ASAH1*$^{-/-}$ HeLa cells, corresponding to **Fig. 5F (bottom)**.

**Dataset S1 (separate file)**: Generation of CRISPR-edited cell lines for interrogation of lysosomal storage disease gene function analysis. This file contains gRNA sequences as well as allele sequencing results for all edits examined. Additionally this file contains annotations of proteins to individual organelles.

**Dataset S2 (separate file)**: nDIA whole cell proteomics of day 50 differentiated iN, iDA of LSD mutants.

**Dataset S3 (separate file)**: nDIA whole cell proteomics of day30 differentiated iN, iDA of select LSD mutants.

**Dataset S4 (separate file)**. Baseline and altered protein-protein interactions.

**Dataset S5 (separate file):** TMT-based analysis of LysoIP sample proteomics of untagged, Control and *ASAH1*$^{-/-}$ HeLa cells.

**Dataset S6 (separate file)**: Label free lipidomics of whole cell and LysoIP of untagged, Control and *ASAH1*$^{-/-}$ HeLa cells.
